# Elongation factor ELOF1 drives transcription-coupled repair and prevents genome instability

**DOI:** 10.1101/2021.05.11.443558

**Authors:** Marit E Geijer, Di Zhou, Kathiresan Selvam, Barbara Steurer, Bastiaan Evers, Chirantani Mukherjee, Simona Cugusi, Marvin van Toorn, Melanie van der Woude, Wenzhi Gong, Roel Janssens, Anja Raams, Joyce HG Lebbink, Bart Geverts, Dalton A Plummer, Karel Bezstarosti, Arjan F Theil, Richard Mitter, Adriaan B Houtsmuller, Wim Vermeulen, Jeroen AA Demmers, Shisheng Li, Hannes Lans, René Bernards, Jesper Q Svejstrup, Arnab Ray Chaudhuri, John J Wyrick, Jurgen A Marteijn

## Abstract

Correct transcription is crucial for life. However, DNA damage severely impedes elongating RNA Polymerase II (Pol II), causing transcription inhibition and transcription-replication conflicts. Cells are equipped with intricate mechanisms to counteract the severe consequence of these transcription-blocking lesions (TBLs). However, the exact mechanism and factors involved remain largely unknown. Here, using a genome-wide CRISPR/cas9 screen, we identified elongation factor ELOF1 as an important new factor in the transcription stress response upon DNA damage. We show that ELOF1 has an evolutionary conserved role in Transcription-Coupled Nucleotide Excision Repair (TC-NER), where it promotes recruitment of the TC-NER factors UVSSA and TFIIH to efficiently repair TBLs and resume transcription. Additionally, ELOF1 modulates transcription to protect cells from transcription-mediated replication stress, thereby preserving genome stability. Thus, ELOF1 protects the transcription machinery from DNA damage by two distinct mechanisms.

## Main Text

Faithful transcription is essential for proper cell function. However, transcription is continuously threatened by DNA damaging agents, which induces transcription-blocking lesions (TBLs) that strongly impede or completely block forward progression of RNA polymerase II (Pol II) ^1,2^. Impeded transcription elongation by DNA damage can affect transcription fidelity or result in complete absence of newly synthesized mRNA transcripts ^3,4^. This can result in severe cellular dysfunction, senescence and cell death, consequently contributing to aging ^5–7^. Furthermore, prolonged stalling of Pol II at TBLs can form physical road blocks for the replication machinery, thereby giving rise to transcription-replication conflicts. These conflicts are detrimental for cells since they can lead to genome instability and onset of cancer ^7–10^. Cells are equipped with an intricately regulated cellular response to overcome the highly toxic consequences of TBLs. This transcription stress response includes repair of TBLs and mechanisms to overcome transcription-replication conflicts ^1,8^.

The main mechanism to remove TBLs is transcription-coupled nucleotide excision repair (TC-NER). TC-NER removes a wide spectrum of environmentally or endogenously-induced TBLs, such UV-light-induced lesions or oxidative damage caused by metabolic processes ^1,11^. The biological consequences of TBLs and relevance of the TC-NER pathway are best illustrated by the fact that inactivating mutations in TC-NER genes can cause Cockayne syndrome (CS), which is characterized by photosensitivity, progressive neurodegeneration and premature aging ^12^. The TC-NER initiating factor CSB (ERCC6) is recruited upon Pol II stalling. CSB uses its forward translocating ability to discriminate between lesion-stalled and other forms of paused Pol II ^13^. When lesion-stalled Pol II is recognized, TC-NER complex assembly is continued by recruitment of CSA (ERCC8) ^14–16^, which is part of a Cullin 4-RING E3 ubiquitin ligase complex (CRL4^CSA^) ^17^, and UVSSA. Interestingly, recently it was shown that also the ubiquitylation of lesion-stalled Pol II plays an important role in the transcription stress response ^18,19^. UVSSA subsequently promotes the recruitment of TFIIH ^18,20^ which forms the core incision complex with XPA and RPA. The incision complex unwinds the DNA, verifies the lesion, and recruits the endonucleases ERCC1/XPF and XPG to excise the TBL ^21^. Repair is finalized by refilling and ligating the gap ^22^, after which transcription can restart. The repair reaction also resolves lesion-stalled Pol II, which helps to lower the frequency of TBL-induced transcription-replication conflicts.

Although several key factors have been identified in the cellular response to DNA damage-induced transcription stress, the exact molecular mechanism to repair TBLs by TC-NER or to avoid collisions of lesion-stalled Pol II with the replication machinery remain largely unknown.

To obtain mechanistic insights in the DNA damage-induced transcription stress response, we performed a genome-wide CRISPR/Cas9 loss-of-function screen to identify novel factors involved in this cellular response. Briefly, fibroblasts were lentivirally transduced with an sgRNA library ^23^ at low multiplicity-of-infection (<0.25). The resulting pool of gene-edited cells was split into two populations. The control group was mock-treated, while in the other group TBLs were induced by exposure to a daily UV-C dose of 6.8 J/m^2^ for 10 consecutive days (Fig. 1A). This UV dose maintained a ∼50% cell confluency throughout the screen (Suppl. Fig 1A). sgRNA abundance was determined by next generation sequencing of PCR-amplified incorporated sgRNAs from the isolated genomic DNA of surviving cell pools ^24^. sgRNA counts from UV-exposed cells were compared to those from untreated cells and negatively selected genes were identified using MAGeCK ^25^ (Fig. 1B, Suppl. Table 1). Gene ontology analysis among the top hits (FDR<0.1), identified many genes involved in the UV-induced DNA damage response (Fig. 1C). These genes included factors involved in translesion synthesis (TLS), like *RAD18* and *POLH* ^26^, and many NER genes including the global genome-NER (GG-NER) damage sensors *DDB2* and *XPC* ^21^. Especially the identification of key TC-NER factors *CSA, CSB* and *UVSSA* underscored the potential of this screen to identify factors involved in the DNA damage-induced transcription stress response ^1^ (Fig. 1B).

**Figure 1.**
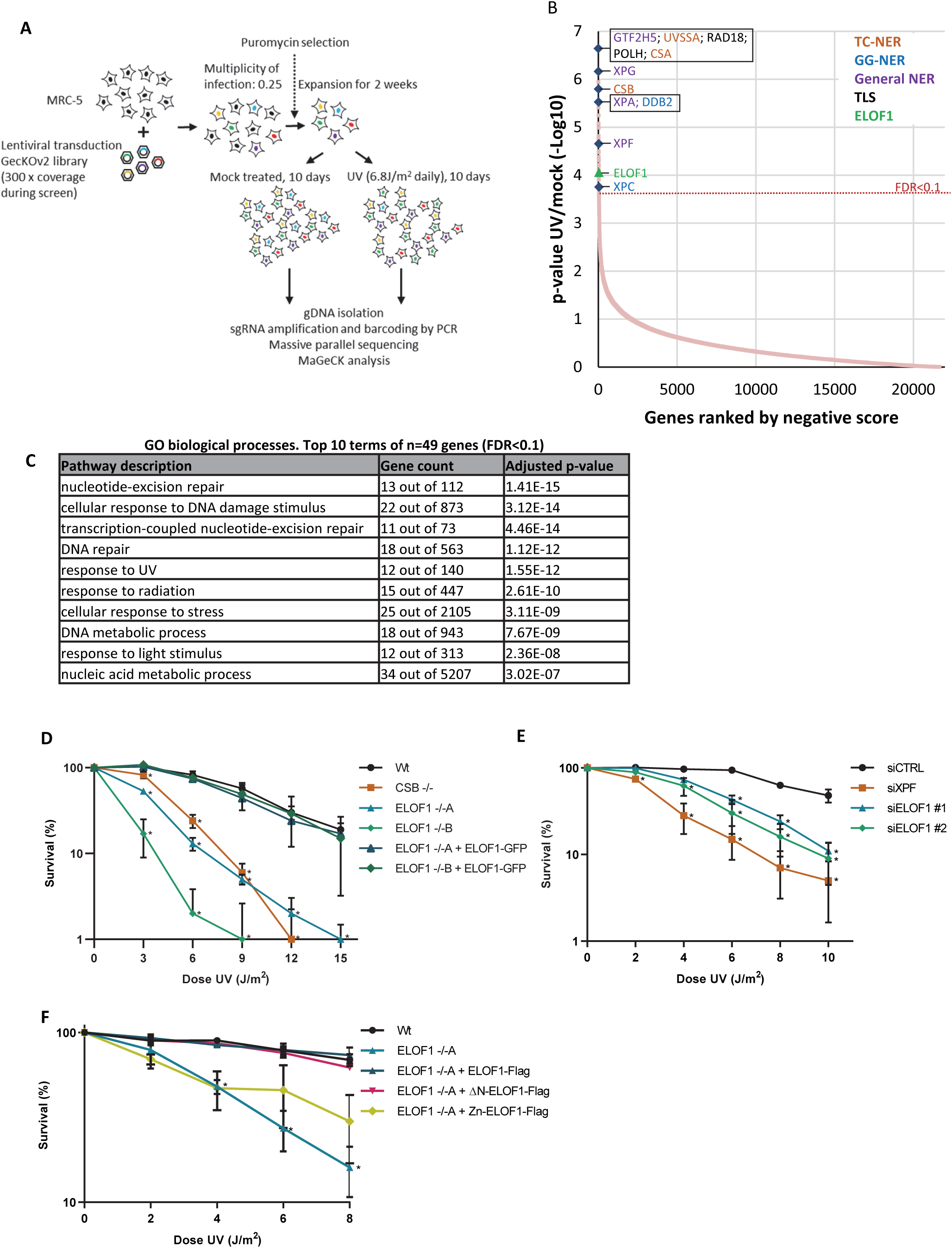
Genome-wide CRISPR/cas9 screen identifies ELOF1 as a novel factor involved in the UV-induced DNA damage response. **(A)** Schematic of the CRISPR/cas9 screen. MRC-5 (SV40) cells infected with a lentiviral sgRNA library were mock-treated or irradiated daily with 6.8 J/m^2^ UV-C for 10 consecutive days. sgRNA abundance was determined by sequencing and UV-sensitive genes were identified by comparing the abundance in UV-irradiated cells over mock-treated cells. The screen was performed in duplicate. **(B)** UV-sensitive genes were ranked based on the gene-based P-value resulting from MaGecK analysis of the change in abundance of sgRNAs in UV-treated over mock-treated. Dotted line indicates FDR=0.1. Genes involved in NER or TLS are color-coded. **(C)** Top 10 enriched GO terms (biological process) identified using g:Profiler of UV-sensitive genes with FDR<0.1 (n=49). **(D)** Relative colony survival of HCT116 wildtype (Wt) cells, indicated knock-out cells (-/-) or rescued cells exposed to the indicated doses of UV-C. **(E)** Relative colony survival of MRC-5 cells transfected with indicated siRNAs following exposure to the indicated doses of UV-C. **(F)** Relative colony survival of HCT116 ELOF1 KO cells with expression of the indicated ELOF1 mutants following exposure to the indicated doses of UV-C. Zn: zinc-finger mutant, ΔN: deletion of N-terminus. Plotted curves represent averages of three independent experiments ± SEM. *P≤0.05.

Interestingly, one of the top hits was Elongation Factor 1 Homolog (*ELOF1*), an evolutionary-conserved small zinc-finger protein (∼10 kDa) ^27^. *ELOF1* was identified in budding yeast in which disruption of its orthologue, *Elf1*, was shown to be synthetic lethal with mutation of genes encoding elongation factors such as *SPT6* and *TFIIS* ^28^. Follow-up studies in yeast revealed that Elf1 interacts with the core Pol II elongation complex as shown by proteomics ^29^, Cryo-EM studies ^30^, and by its presence at gene bodies as shown by ChIP ^28,31^. *In vitro* studies showed that Elf1 binds downstream of Pol II at the DNA entry tunnel and promotes elongation through nucleosomes ^32^. However, its exact function as a transcription elongation factor, especially in mammalian cells, and its role in the DNA damage response has thus far remained unknown.

To validate the sensitivity to UV upon ELOF1 depletion, as determined in our CRISPR/Cas9 screen, we performed clonogenic survival experiments using two independent ELOF1 knockout (KO) cell lines (Suppl. Fig. 1B-F). ELOF1 KO resulted in a severe UV hypersensitivity, even slightly higher than observed in TC-NER-deficient CSB KO cells (Fig. 1D). Similar results were obtained upon siRNA-mediated depletion of ELOF1 (Fig. 1E, Suppl. Fig. 1G,H). Re-expression of ELOF1 in ELOF1 KO cells fully rescued their UV sensitivity, indicating that the observed effects are specific for ELOF1. Although the N-terminal tail of ELOF1 promotes Pol II progression on the nucleosome ^32^, constructs without this N-terminal tail could still rescue the UV sensitivity in ELOF1 KO cells (Fig, 1F, Suppl Fig. 1I). However, the conserved zinc-finger domain of ELOF1 was crucial for survival upon UV-induced DNA damage. Furthermore, photolyase-mediated reversal of UV-induced cyclobutane pyrimidine dimer (CPD) lesions (33) almost completely rescued the UV sensitivity of ELOF1 KO cells, showing that this sensitivity is due to induction of DNA damage and not RNA or protein damage (Suppl. Fig. 1J).

We first tested whether ELOF1 is part of the elongating Pol II complex, as previously observed for Elf1 in yeast ^28,29^. Since we could not obtain antibodies capable of recognizing endogenous ELOF1, we generated homozygous *ELOF1-mScarletI-HA* knock-in (KI) cells to allow detection of endogenously expressed ELOF1 (Suppl. Fig. 2A). In living cells, ELOF1 was localized strictly to the nucleus, excluded from the nucleoli, and showed high level of co-localization with endogenously expressed GFP-tagged RPB1 ^33^, the largest subunit of Pol II (Fig. 2A and Suppl. Fig. 2B,C). Previous live-cell imaging studies on GFP-RPB1 mobility showed that fluorescence recovery after photobleaching (FRAP) experiments are a sensitive way to study Pol II-mediated transcription, as the different steps of the transcription cycle are characterized by kinetically distinct Pol II populations ^33^. Therefore, we compared the mobility of ELOF1 to that of Pol II using FRAP, and observed that it was almost identical in non-treated conditions (Fig. 2B). The large ELOF1 immobilization, which slowly redistributes in time, suggests that the majority of ELOF1 molecules is chromatin-bound, most likely engaged in transcription elongation, similar as observed for Pol II ^33^. The engagement of ELOF1 in transcription elongation was confirmed by its swift chromatin release, as shown by its strong mobilization upon inhibition of transcription initiation with the CDK7 inhibitor THZ1 ^34^, or inhibition of release of promoter-paused Pol II into the gene body by the CDK9 inhibitor Flavopiridol ^35^ (Fig 2B). This almost complete mobilization upon transcription inhibition suggests that ELOF1 is exclusively involved in transcription-related processes. Furthermore, this highly similar dynamic behavior, also upon transcription inhibition, suggests that ELOF1 is closely associated with Pol II. Interestingly, treatment with the DNA intercalator actinomycin D ^36^ resulted in a severe immobilization of Pol II, while ELOF1 was only transiently immobilized. This suggests that ELOF1 can still dissociate from actinomycin D-stalled Pol II complexes while RPB1 remains trapped on the DNA.

**Figure 2.**
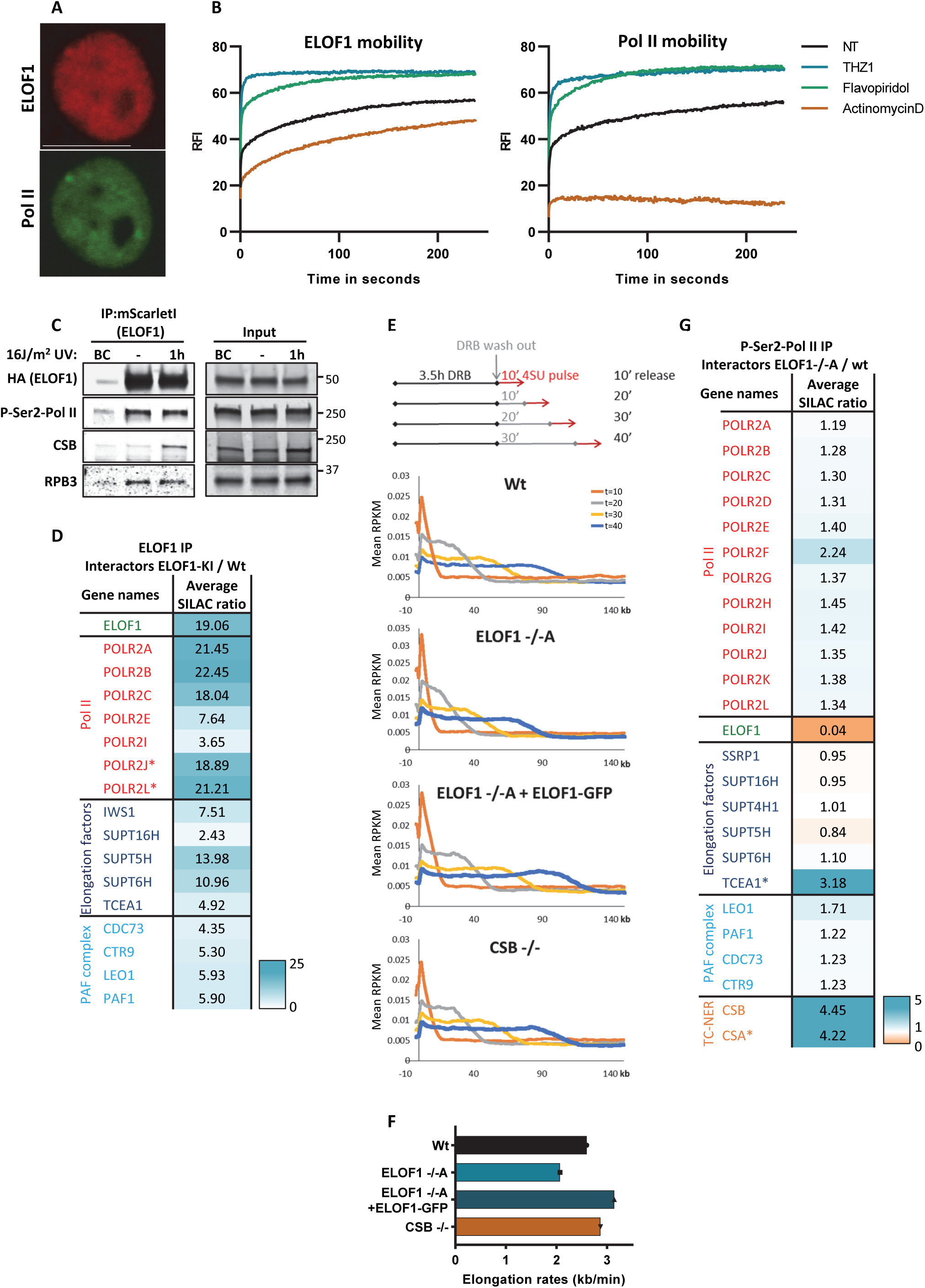
ELOF1 is part of the elongating Pol II complex. **(A)** Co-localization of ELOF1 and Pol II in HCT116 cells with ELOF1-mScarletI-HA and GFP-RPB1 during live-cell imaging. Scale bar: 10 µm. **(B)** Fluorescence recovery after photobleaching (FRAP) analysis of endogenously expressed ELOF1-mScarletI (Left) and GFP-RPB1 (Right). Cells were mock-treated (NT) or inhibited at different steps of the transcription cycle using indicated inhibitors. Relative Fluorescence Intensity (RFI) was measured over time, background-corrected, and normalized to pre-bleach fluorescence intensity. n≥20 for *ELOF1*-KI and n≥8 for *RPB1*-KI cells. **(C)** Immunoprecipitation of ELOF1 using RFP beads in *ELOF1*-KI cells followed by immunoblotting for indicated proteins. Cells were harvested 1 hour after mock treatment or irradiation with 16 J/m^2^ UV-C. BC: binding control. **(D)** Interaction heat map of the SILAC ratios of ELOF1-interacting proteins as determined by quantitative interaction proteomics following HA-IP of ELOF1. Average SILAC ratios of duplicate experiments are plotted and represent ELOF1-interactors relative to empty beads. SILAC ratio >1 indicate increase in interaction. * indicates proteins quantified in one experiment. **(E)** Top panel: Schematic of DRB/TT_chem_-seq to measure Pol II elongation rates. Bottom panel: Metagene profiles of DRB/TT_chem_-seq in HCT116 Wt or indicated KO (-/-) cells, with ELOF1 re-expression where indicated, 10, 20, 30, or 40 minutes after DRB release. **(F)** Average elongation rates as determined by DRB/TT_chem_-seq. **(G)** Interaction heat map based on the SILAC ratios as determined by quantitative interaction proteomics of P-Ser2-modified Pol II-interacting proteins in ELOF1 -/-A cells relative to Wt cells. Average SILAC ratios of duplicate experiments are plotted. * indicates proteins quantified in one experiment. SILAC ratios <1 indicate loss of interaction, >1 indicate increase in interaction.

To further investigate whether ELOF1 is part of the elongating Pol II complex we immunoprecipitated (IP) ELOF1 and detected its interaction with the RPB1 and RPB3 subunits of Pol II (Fig. 2C). The interaction of ELOF1 with P-Ser2-modified RPB1, which primarily marks productively elongating Pol II, indicates that ELOF1 is present in the elongating Pol II complex. The reciprocal IP of P-Ser2-modified Pol II confirmed that ELOF1 interacts with elongating Pol II (Suppl. Fig. 2D). Moreover, SILAC-based interaction proteomics of endogenously expressed GFP-RPB1 ^33^ identified ELOF1 as a genuine Pol II interactor with similar SILAC ratios as other elongation factors (Suppl. Fig. 2E). To obtain a complete overview of ELOF1 protein interactions, we performed SILAC-based interaction proteomics for ELOF1, revealing high SILAC ratios for many Pol II subunits and elongation factors including TFIIS, SPT6, SPT5 and the PAF complex ^37,38^ (Fig. 2D and Suppl. Table 2). Gene ontology analysis of the most enriched ELOF1-interactors showed specific involvement in transcription-related processes (Suppl. Fig. 2F). Of note, the ELOF1-Pol II interaction did not change upon UV-induced DNA damage, in contrast to the interaction with CSB ^39^ (Fig. 2C and Suppl. Fig. 2D). Together, these live-cell imaging and interaction data indicated that ELOF1 is an integral component of the transcription elongation complex, independent of DNA damage.

Next, we tested whether ELOF1 acts as a transcription elongation factor by determining its effect on Pol II elongation rates. Therefore, we performed DRB/TT_chem_-seq ^40^, in which nascent RNA is labeled with 4SU to determine the Pol II position in a gene body at different time points after its release from the promoter by DRB washout (Fig. 2E). Single gene profiles (Suppl. Fig. 3A) and metagene analysis (Fig. 2E) showed that ELOF1 KO resulted in a clear decrease in elongation rate, while ∼6-fold overexpression of ELOF1 (Suppl. Fig. 1E) resulted in increased elongation rate. Based on metagene analysis, an average decrease in elongation speed from 2.6 kb/min to 2.0 kb/min was observed for ELOF1 KO cells, while an increase to 3.1 kb/min was observed after ELOF1 overexpression (Fig. 2F). In line with this reduced elongation rate, the overall nascent RNA synthesis was also reduced upon ELOF1 depletion (Suppl. Fig. 3B-D). In contrast, loss of CSB had no obvious effect on Pol II elongation rate.

To identify the mechanism for the reduction in elongation speed after loss of ELOF1, we compared the composition of the elongation complex with and without ELOF1. Endogenous P-Ser2-modified Pol II was isolated, and differences in the Pol II interactome were detected using SILAC-based proteomics. Absence of ELOF1 did not affect the presence of the core Pol II subunits or the majority of elongation factors in the elongation complex (Fig. 2G, Suppl. Table 2). For example, presence of the SPT4/5 dimer, which interacts genetically and biochemically with yeast ELOF1 ^28,32^, was not changed in the elongation complex. Interestingly, the biggest change in complex composition was found for CSA, CSB and TFIIS, each having a 3- to 4-fold increased interaction with Pol II without ELOF1 (Fig. 2G). As CSB recognizes stalled and paused Pol II complexes, for example at DNA lesions or natural pause sites ^13^, the increase in CSB binding might indicate that the forward translocation of Pol II is more frequently perturbed in the absence of ELOF1. Such perturbation can induce Pol II backtracking that is recognized by TFIIS to stimulate subsequent transcript cleavage ^41^ to allow continued forward translocation. In line with such a model, we observed increased TFIIS binding to elongating Pol II in ELOF1 KO cells (Fig. 2G). In addition, depletion of TFIIS gave rise to synthetic lethality with ELOF1 KO (Suppl. Fig. 3E) as was previously also observed in yeast ^28^.

After having established that ELOF1 is a *bona fide* elongation factor, we studied its role in the DNA damage response. Since ELOF1 was shown to be an integral part of the transcription elongation machinery, and ELOF1 KO cells are sensitive for UV-induced DNA damage, which is a potent inhibitor of transcription, we tested whether ELOF1 is needed for recovery of transcription after UV irradiation by quantifying nascent transcription levels by EU incorporation ^42^. Transcription was severely reduced 2 hours after UV damage but fully recovered in Wt cells after 18 hours (Fig. 3A and Suppl. Fig. 4A). Strikingly, the transcription recovery was completely abolished in ELOF1 KO cells, as in TC-NER deficient CSA KO cells, but could be rescued by re-expression of ELOF1. Similar results were obtained using siRNA-mediated ELOF1 knockdown (Suppl. Fig. 4B,C). This indicates that ELOF1 either has a function in the removal of TBLs, or in the restart of transcription. To distinguish between both possibilities, we measured TC-NER activity by quantifying the gap-filling synthesis using EdU incorporation in non-replicating GG-NER deficient cells ^43^. Like CSB depletion, loss of ELOF1 severely inhibited the TC-NER activity indicating that ELOF1 has a crucial function in TC-NER (Fig. 3B, Suppl. Fig. 4D-E). The function of ELOF1 was restricted to the TC-NER sub-pathway, since the gap-filling synthesis in GG-NER proficient cells was not affected (Fig. 3C, Suppl. Fig. 4F-G). Together, this shows that ELOF1 is important for removing UV-induced lesions by TC-NER to subsequently promote transcription recovery.

**Figure 3.**
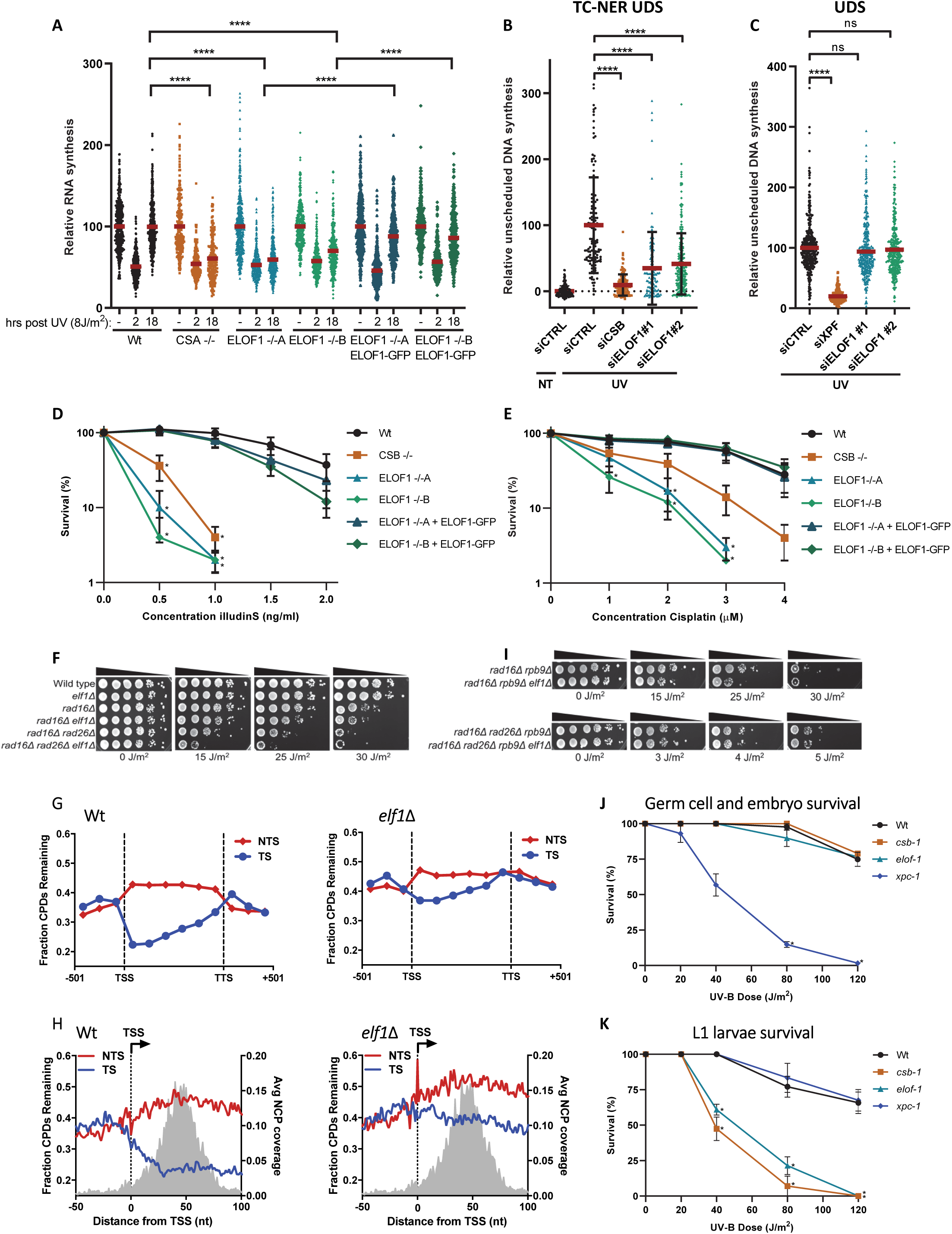
ELOF1 is an evolutionary-conserved core TC-NER factor. **(A)** Transcription restart after UV damage as determined by relative EU incorporation in the indicated HCT116 Wt and KO (-/-) cells, with ELOF1 re-expression where indicated, at the indicated time points after UV-C (8 J/m^2^). Relative integrated density normalized to mock-treated levels and set to 100%. Red lines indicate average integrated density ± SEM. n≥300 cells from at least three independent experiments. **(B)** TC-NER-specific UDS as determined by relative EdU incorporation in XP186LV fibroblasts (XP-C) transfected with indicated siRNAs following UV-C-irradiation (7 hours, 8 J/m^2^). n≥100 cells from two independent experiments. **(C)** Relative levels of EdU incorporation in C5RO (hTert) cells transfected with indicated siRNAs, following UV-C-irradiation (3 hours, 16 J/m^2^). n≥200 cells from at least two independent experiments **(D+E)** Relative colony survival of the indicated HCT116 Wt and KO (-/-) cells, with ELOF1 re-expression where indicated, upon a 24-hour exposure to the indicated concentrations of illudinS **(D)** or Cisplatin **(E).** Plotted curves represent average of at least three independent experiments ± SEM. **(F)** Indicated mutant yeast strains were serially 10-fold diluted, spotted, and exposed to the indicated UV-C doses. **(G)** CPD-seq analysis of Wt (left) and *elf1Δ* mutant (right) yeast showing the average fraction of unrepaired CPDs remaining on the transcribed strand (TS) and non-transcribed strand (NTS) for ∼4500 yeast genes following 2-hour repair relative to no repair. Each gene was divided in 6 equally-sized bins. Repair in flanking DNA upstream of the transcription start site (TSS) and downstream of the transcription termination site (TTS) is also depicted. **(H)** Close-up of CPD-seq repair data near the TSS in Wt (left) and *elf1Δ* mutant (right) cells. Nucleosome positioning data is shown for reference. **(I)** Indicated mutant yeast strains were serially 10-fold diluted, spotted, and exposed to the indicated UV-C doses. **(J)** *C. elegans* germ cell and embryo UV survival assay, measuring GG-NER activity, of wild type, *csb-1, xpc-1,* and *elof-1* animals. The percentages of hatched eggs (survival) are plotted against the applied UV-B doses. The mean survival of two replicate experiments each performed in quintuple is depicted. **(K)** L1 larvae UV survival assay, measuring TC-NER activity, of wildtype, *csb-1*, *xpc-1* and *elof-1* animals. The percentages of animals that developed beyond the L2 stage (survival) are plotted against the applied UV-B doses. The mean survival of three replicate experiments each performed in quintuple is depicted. Error bars represent the SEM. *p≤0.05, ****p≤0.0001.

Next, we tested the sensitivity of ELOF1 KO cells to other types of DNA damage. Interestingly, ELOF1 KO, like CSB KO, resulted in a severe sensitivity to a wide spectrum of genotoxins that cause TBLs, including Illudin S ^44^, Cisplatin ^45^, Camptothecin ^46^ and oxidative lesions ^47^ (Fig. 3D-E and Suppl. Fig. 5A-C). However, ELOF1 KO cells were not sensitive to replication stress induced by hydroxyurea (Suppl. Fig. 5D). Importantly, sensitivity to TBLs is generally not observed after depletion of elongation factors since these were not among the top hits of our CRISPR/Cas9 screen for UV-sensitive genes (Suppl. Table 1). In addition, transient knockdown of the core elongation factors SPT4 and SPT5 did not increase UV sensitivity, although this caused a comparable reduction in RNA synthesis, similar as depletion of ELOF1 (Suppl. Fig. 3B-D, 5E).

As *ELOF1* is highly conserved from archaea to mammals ^27^, we tested whether the *ELOF1* orthologues in *Saccharomyces cerevisiae* and *Caenorhabditis elegans* are also involved in repairing TBLs. Similar to mutations in *RAD26,* the budding yeast ortholog of *CSB*, inactivation of *ELF1* (*elf1Δ*) had no effect on the UV sensitivity (Suppl. Figure 6A), which can be explained by the highly efficient GG-NER machinery in budding yeast ^48^. To specifically study the effect of *elf1Δ* in TC-NER, we tested the effect of its inactivation on UV survival in GG-NER-deficient *RAD16* mutants (*rad16Δ*). This showed a clearly increased sensitivity to UV for both the *elf1Δ* and the *rad26Δ* mutants, suggesting that Elf1 is involved in TC-NER (Fig. 3F).

To determine if the increased UV sensitivity in the *elf1Δ* mutant is caused by a TC-NER-defect, we analyzed CPD repair profiles in the transcribed strand (TS) and non-transcribed strand (NTS) of yeast genes 2 hours after UV using high-resolution CPD-sequencing ^49^ (Suppl. Fig. 6B). This analysis showed that in the *elf1Δ* mutant, GG-NER-mediated repair in the NTS was hardly affected. However, TC-NER-mediated repair in the TS was severely compromised, as shown by meta-analysis of ∼4500 genes (Fig. 3G) and by individual genes (Suppl. Fig. 6C). The global repair rate in *elf1Δ* was hardly affected (Supp. Fig. 6D), which is in agreement with a TC-NER-specific effect that only happens in the TS of active genes. Although Elf1 was described to stimulate Pol II progression on the nucleosome ^32^, no nucleosome-dependent difference in TC-NER efficiency was detected in the TS in *elf1Δ* mutants (Suppl. Fig. 6E). Moreover, deletion of the N-terminus of ELOF1, which is involved in transcription processivity at nucleosomes ^32^, had no effect on the UV-survival (Fig. 1F).

Strikingly, the *elf1Δ rad26Δ* double mutant showed an even higher UV sensitivity than the *elf1Δ* and *rad26Δ* single mutants in a *rad16Δ* background, indicating that Elf1 has functions in the UV-induced DNA damage response independent of Rad26 (Fig. 3F). Close-ups of the CPD sequencing data showed that repair in the *elf1Δ* mutant is also compromised immediately downstream of the transcription start site (TSS) (Fig. 3H). This genomic region can be repaired in a Rad26-independent manner ^50^ by a Rpb9-mediated transcription-coupled repair mechanism ^51^. This may suggest that Elf1 also functions in Rpb9-mediated repair, independent of Rad26 (Fig. 3F). Indeed, *elf1*Δ enhances the UV sensitivity in *rad16Δrpb9Δ* mutants, but not in a *rad16Δrpb9Δrad26* mutant (Fig 3I and Suppl Fig. 6F), indicating that Elf1 is involved in both Rad26-dependent and -independent repair. This was confirmed by the finding that deletion of *ELF1* in both *rad16Δrad26Δ* and *rad16Δrpb9Δ* mutants resulted in reduced TC-NER (Suppl. Fig. 6G,H).

To study the role of ELOF1 in a multi-cellular model organism, we made use of the conservation of *ELOF1* in *C. elegans*. We assayed UV-survival of mutant germ and early embryonic cells, which predominantly depends on GG-NER, and of post-mitotic first-stage larvae, which mainly depends on TC-NER ^52^. In contrast to inactivation of the GG-NER factor *xpc-1,* did inactivation of *elof-1* not increase UV sensitivity of germ and embryonic cells (Fig. 3J and Suppl. Fig. 6I). However, *elof-1* mutant animals showed a strong UV sensitivity in the first larval stage, similar to the TC-NER-deficient *csb-1* animals (Fig. 3K).

Together these data indicate that ELOF1 is an important and highly evolutionary-conserved repair factor, specifically involved in repair of DNA damage in transcribed strands of active genes (Fig. 3). As ELOF1 is an integral part of the elongation complex (Fig. 2), its depletion will most likely affect Pol II forward translocation upon encountering TBLs. To test this, we used GFP-RPB1 KI cells to study Pol II mobility by FRAP, which provides quantitative information on Pol II elongation rates and fraction sizes of elongating and promoter-bound Pol II, i.e. initiating and promotor-paused Pol II ^33^. UV-induced DNA damage resulted in an increased Pol II immobilization, especially of the long-bound fraction, as evident from the reduced slope of the FRAP curve at time points >100 sec (Fig. 4A), which mainly represents dynamics of elongating Pol II ^33^. Monte-Carlo-based modeling ^53^ of these FRAP data revealed an increase in the fraction size and residence time of elongating Pol II. This indicates that UV-exposure resulted in more elongating Pol II transcribing with a lower average elongation rate (Fig 4B), most likely caused by Pol II stalling at TBLs ^54,55^. Interestingly, the long-bound Pol II fraction after UV was further immobilized upon knockdown of ELOF1, to a similar extent as after depletion of CSB. Monte-Carlo-based modeling of these FRAP data showed an approximately 30% increase of the average residence time of elongating Pol II, suggesting that Pol II stalling at lesions is prolonged in the absence of ELOF1 (Fig. 4A-B, Suppl. Fig. 7C). Similar results were obtained by Pol II ChIP-seq experiments (van der Weegen *et al.* submitted back-to-back). ELOF1 knockdown also resulted in an increased residence time of elongating Pol II in unperturbed conditions, indicative of a reduced elongation rate (Fig. Suppl. 7A-B), in line with our DRB/TT_chem_-seq data (Fig 2E-F).

**Figure 4.**
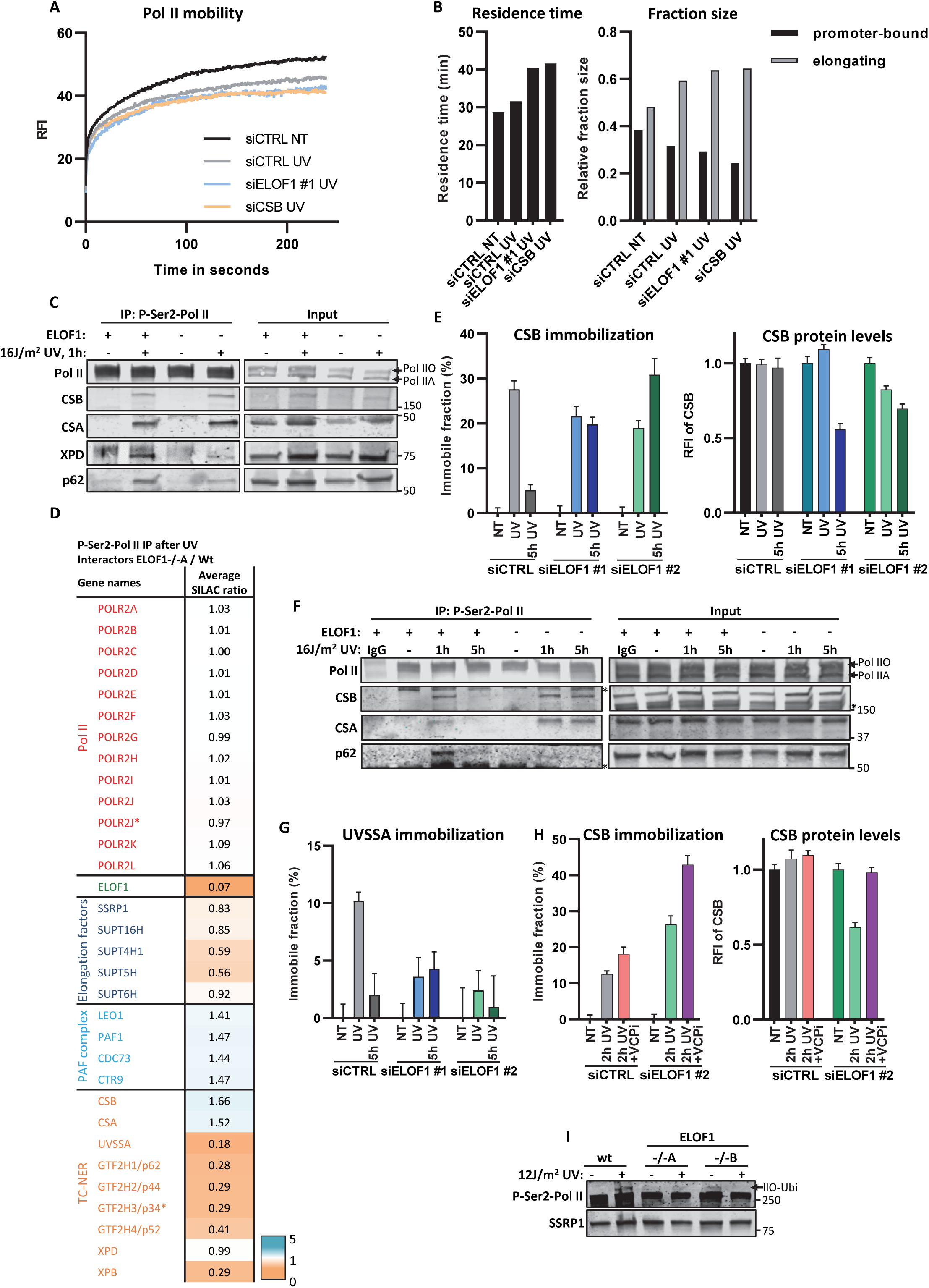
ELOF1 is crucial for proper TC-NER complex assembly. **(A)** FRAP analysis of Pol II mobility in MRC-5 *GFP-RPB1* KI cells after depletion of indicated factors in untreated cells (NT) or directly after UV induction (UV, 12 J/m^2^). Relative Fluorescence Intensity (RFI) was measured over time, background-corrected, and normalized to pre-bleach fluorescence intensity. n≥17 cells. **(B)** Left panel: residence time of the elongating Pol II fraction. Right panel: relative fraction sizes of promoter-bound or elongating Pol II as determined by Monte-Carlo-based modeling based on the RPB1 mobility shown in (A). **(C)** Immunoprecipitation of P-Ser2-modified Pol II in Wt and ELOF -/-A cells followed by immunoblotting for indicated proteins. Cells were harvested 1 hour after mock treatment or irradiation with 16 J/m^2^ UV-C. **(D)** Interaction heat map based on the SILAC ratios as determined by quantitative interaction proteomics of UV-specific Pol II-interacting proteins in ELOF1 -/-A cells relative to Wt cells. Average SILAC ratios of duplicate experiments are plotted. SILAC ratios <1 indicate loss of interaction, >1 indicate increase in interaction. * indicates proteins quantified in one experiment. **(E)** Left panel: Relative immobile fraction of CSB in *CSB-mScarletI* KI cells transfected with indicated siRNAs directly (UV) or 5 hours after UV-C irradiation (5h UV, 4 J/m^2^) as determined by FRAP analysis (Suppl. Fig. S8E). Right panel: Relative fluorescence intensity of CSB-mScarletI in *CSB*-KI cells transfected with indicated siRNAs as determined by live-cell imaging. Plotted values represent mean ± SEM and are normalized to mock-treated. n≥9 cells. **(F)** Immunoprecipitation of P-Ser2-modified Pol II in Wt and ELOF -/-A cells 1 hour or 5 hours after UV-C (16 J/m^2^) irradiation followed by immunoblotting for indicated proteins. IgG was used as binding control. *a-specific band. **(G)** Same as left panel of E but for *UVSSA-mScarletI* KI cells (Suppl. Fig. 8F-G). n≥16 cells. **(H)** Relative immobile fraction (left panel) or relative fluorescence intensity (right panel) of CSB-mScarletI in *CSB*-KI cells transfected with indicated siRNAs 2 hours after UV-C irradiation (4 J/m^2^) as determined by FRAP analysis (Suppl. Fig. 8H). VCPi: treatment with VCP inhibitor. Plotted values represent mean ± SEM and are normalized to mock-treated. n≥10 cells. **(I)** Immunoblot of chromatin fraction of indicated HCT116 Wt or ELOF1 KO cells 1 hour after 12 J/m^2^ UV-C or mock treatment. SSRP1 is shown as loading control.

Since Pol II elongation was slowed down upon DNA damage induction in the absence of ELOF1, we immunoprecipitated elongating Pol II after UV to study whether specific reaction steps of TC-NER were compromised. In ELOF1 KO cells, TC-NER initiating factors CSA and CSB were still properly bound to lesion-stalled Pol II. However, the UV-induced Pol II interaction with TFIIH complex subunits XPD and p62 was strongly reduced (Fig. 4C), which was not a consequence of TFIIH degradation (Suppl. Fig. 7D). To obtain a more unbiased overview of the effects of ELOF1 KO on the DNA damage-induced Pol II interactome, we performed SILAC-based interaction proteomics on the elongation complex after UV-induced DNA damage in presence or absence of ELOF1. The interaction of most elongation factors with Pol II was not affected by the absence of ELOF1 (Fig.4D and Suppl. Fig.7E). Interestingly, while CSA and CSB could still bind to Pol II in the absence of ELOF1, the proteins most affected in their Pol II binding were UVSSA and TFIIH subunits.

Since UVSSA plays a crucial role in the recruitment of TFIIH to lesion-stalled Pol II ^18,20,56^, the decreased binding of UVSSA is most likely the cause of reduced TFIIH recruitment, and explains the observed TC-NER defects. To confirm these results, we generated *CSB* and *UVSSA* knock-in cells (Suppl. Fig. 8A,B), expressing mScarletI-tagged CSB and UVSSA proteins from their endogenous locus, allowing direct analysis of their quantity and mobility in living cells. TBL-induced immobilization of these TC-NER factors, as determined by FRAP ^39,57^, is an accurate measure for their involvement in TC-NER, as shown by their UV-induced immobilization in a transcription-dependent manner (Suppl. Fig. 8C,D). In line with the IP experiments, did ELOF1 depletion not affect the CSB immobilization 1 hour after UV. CSB remained immobilized up to at least 5 hours after UV (Fig. 4E and Suppl. Fig. 8E). This prolonged binding of CSB to stalled Pol II upon UV was confirmed by IP experiments, which also showed prolonged binding of CSA (Fig. 4F). These observations are in line with a model in which TBLs cannot be removed because of the TC-NER defect caused by ELOF1-deficiency, which will result in prolonged binding of CSB and CSA to lesion-stalled Pol II. In contrast, UVSSA immobilization upon UV damage was severely reduced after ELOF1 depletion (Fig. 4G and Suppl. Fig. 8F-G), further indicating that ELOF1 plays a crucial role in the recruitment of UVSSA to lesion-stalled Pol II. UVSSA recruits the deubiquitylating enzyme USP7, which protects CSB from proteasomal degradation mediated by the ubiquitin-selective segregase VCP/p97 ^57–59^. In line with this, we observed that reduced UVSSA recruitment upon ELOF1 depletion, and consequently of USP7, resulted in a UV-induced ∼40% decrease of overall CSB levels (Fig. 4E) by VCP-mediated proteasomal degradation (Fig. 4H right panel). Interestingly, FRAP analysis showed an even stronger CSB immobilization upon TBL induction and VCP inhibition in ELOF1-depleted cells (Fig. 4H and Suppl. Fig. 8H), suggesting that chromatin-bound CSB is degraded in the absence of ELOF1, most likely when bound to lesion-stalled Pol II. Therefore, the increase in CSB immobilization upon UV-exposure confirms that in the absence of ELOF1, a larger Pol II fraction remains stalled at the lesion.

Recently the ubiquitylation of a single lysine mutation in RPB1 (K1268) was described to be an important event in the transcription stress response ^18,19^. Therefore, we tested whether ELOF1 is involved in the UV-induced ubiquitylation of Pol II by means of a slower migrating P-Ser2-modified RPB1 band ^18^. Interestingly, ELOF1 KO almost completely abolished the UV-induced RPB1 ubiquitylation (Fig. 4I), to the same extent as CSB KO or inhibiting NEDD8-conjugating enzyme NAE1, which controls the activity of CRL complexes ^18,19^ (Suppl. Fig. 8I). Similar results were obtained using siRNA-mediated depletion of ELOF1 (Suppl. Fig. 8J).

Together, our results demonstrate that the presence of ELOF1 in the lesion-stalled Pol II complex is an important determinant for proper Pol II ubiquitylation and for correct assembly of the TC-NER complex. The observed TC-NER defect explains the severe sensitivity of ELOF1 KO cells to different TBLs (Fig. 1D,E, 3D,E and Suppl. Fig. 5A-C). Strikingly, while testing the sensitivity of ELOF1 KO cells for a wide spectrum of DNA lesions, we observed that ELOF1 KO cells were also sensitive to the DNA crosslinker Mitomycin C (MMC) (Fig. 5C). Interestingly, CSB KO cells were not sensitive to MMC suggesting that ELOF1 has an additional function in the DNA damage response, besides canonical TC-NER. The prolonged transcription block in ELOF1 KO cells upon MMC exposure (Suppl. Fig. 9A), which was not observed in CSB KO cells, suggests that this additional role for ELOF1 is also linked to transcription. To investigate this additional function of ELOF1, we depleted ELOF1 in TC-NER-deficient CSB KO or NER-deficient XPA KO cells and strikingly observed that this resulted in increased UV sensitivity (Fig. 5B). Interestingly, CSB has also additional functions to ELOF1 in the response to UV-induced damage (Suppl. Fig. 9B). The additional role of ELOF1 to TC-NER was further confirmed in CS patient cells characterized by inactivating mutations in CSA (CS-A), in which knockdown of ELOF1 also resulted in additional UV sensitivity (Fig. 5C). Remarkably, this additive effect was completely absent in non-cycling CS-A cells (Fig. 5D), indicating that it is dependent on cell proliferation.

**Figure 5.**
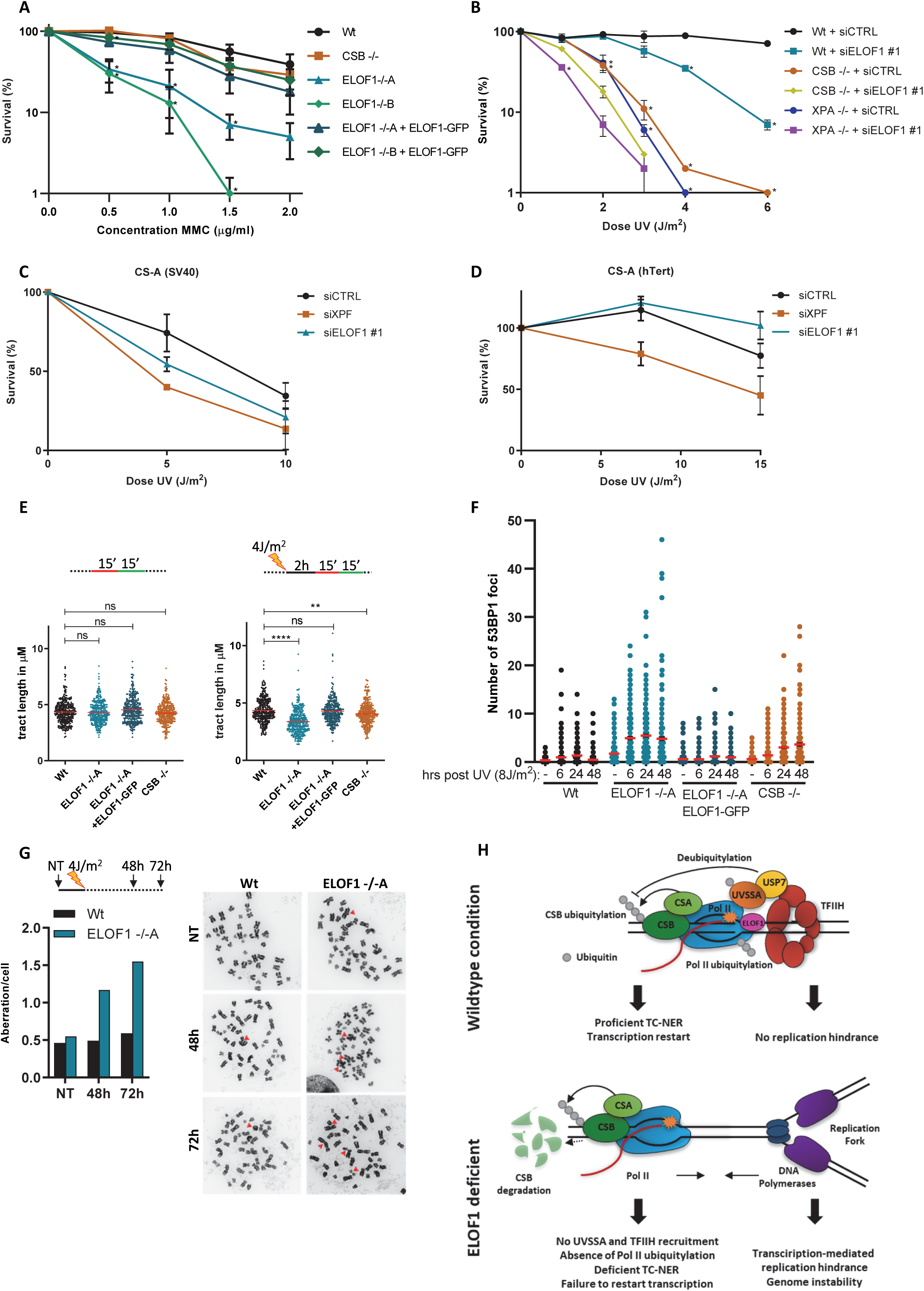
ELOF1 is important for preventing genome instability in addition to its function in TC-NER. **(A)** Relative colony survival of indicated HCT116 Wt and KO (-/-) cells, with ELOF1 re-expression where indicated, upon a 1-hour exposure to the indicated concentrations of mitomycin C. Plotted curves represent averages of three independent experiments ± SEM. **(B)** Relative colony survival of MRC-5 Wt or indicated KO (-/-) cell lines, transfected with indicated siRNAs following exposure to the indicated doses of UV. Plotted curves represent averages of at least two independent experiments ± SEM. *P≤0.05. **(C+D)** Viability of replicating CS-A (SV40, C) or non-replicating primary CS-A cells (hTert, D) following exposure to the indicated UV-C doses as determined by AlamarBlue staining. Plotted curves represent averages of at least two experiments ± SEM. **(E)** Top panel: Schematic of experimental conditions for fork progression in indicated cell lines labeled with CldU (red) for 15 min followed by IdU (green) for 15 min as indicated. Bottom panel: Fork progression measured by tract lengths of CldU (red) in micrometers (μM) is depicted for HCT116 Wt and KO (-/-) cells, with ELOF1 re-expression where indicated, in untreated conditions (left) or 2 hours after 4 J/m^2^ UV-C (right). n≥300 tracts from three independent experiments. **(F)** Number of 53BP1 foci in HCT116 Wt and KO (-/-) cells, with ELOF1 re-expression where indicated, in untreated conditions or the indicated time after UV-C (8 J/m^2^). Red lines indicate average number of foci ± SEM. n≥200 cells from two independent experiments. **(G)** Left panel: Quantitation of chromosomal aberrations per cell in HCT116 Wt and ELOF1 -/-A cells 48 or 72 hours after irradiation with 4 J/m^2^ UV-C or mock treatment (NT). At least 60 metaphases were analyzed. Right panel: Representative images of metaphase spreads. Arrows indicate chromosomal aberrations. **(H)** Model showing function of ELOF1. Top panel: wildtype conditions: ELOF1 is an integral part of the elongation complex and binds near the DNA entry tunnel and ubiquitylation site of Pol II to promote TC-NER and subsequent transcription restart. Cells do not have replication problems. Bottom panel: in the absence of ELOF1 are CSA and CSB still recruited to lesion-stalled Pol II, however, UVSSA, TFIIH, and Pol II ubiquitylation are absent. The incomplete assembly of the TC-NER complex prevents functional TC-NER and subsequent transcription restart. In addition, there is an increase in transcription-mediated replication stress leading to genome instability. *p≤0.05, **≤0.01, ****p≤0.0001.

This replication-dependent sensitivity, together with the specific role of ELOF1 in transcription (Fig. 2), its additive effect to TC-NER (Fig. 5A and Suppl. Fig. 8A), and the prolonged Pol II stalling upon ELOF1 knockdown (Fig. 4A,B), opened the possibility that lesion-stalled Pol II collides with incoming replication forks in the absence of ELOF1, thereby causing transcription-replication conflicts. Therefore, we investigated the impact of ELOF1 KO on DNA replication by analyzing the progression rates of individual replication forks by sequentially labelling cells with CldU and IdU. Tract length analysis revealed no significant difference in replication fork progression upon ELOF1 KO in unperturbed conditions, indicating that ELOF1 has no role in fork progression (Fig. 5E). However, 2 hours after UV, the tract length was significantly decreased in ELOF1 KO cells compared to Wt and ELOF1-complemented cells. Also, in CSB KO cells a small effect on fork progression was observed, however not to the same extent as in ELOF1 KO cells. This suggests that loss of the elongation factor ELOF1 results in replication problems upon induction of TBLs, likely due to the transcription-mediated replication blockage. Transcription-replication conflicts have previously been shown to result in under-replicated DNA, which may cause DSBs upon mitotic progression and subsequently give rise to genome instability ^8,60^. In line with this hypothesis, we observed a more pronounced increase in 53BP1-foci upon UV irradiation in ELOF1 KO cells compared to Wt or CSB KO cells (Fig. 5F, Suppl. Fig. 9C). As replication-interference and under-replicated DNA are important drivers of chromosomal aberrations, we assessed this in ELOF1 KO cells. This clearly resulted in an increased number of chromosomal aberrations in ELOF1 KO cells compared to Wt cells (Fig 5G).

We unveiled an important role for ELOF1 in the cellular response to DNA damage-induced transcription stress by two independent mechanisms: promoting TC-NER and reducing transcription-mediated replication hindrance (Fig. 5H). First, ELOF1 is crucial for TC-NER. Interestingly, while the interaction of most TC-NER factors with elongating Pol II is strongly increased upon DNA damage ^1,14,39, 56–58^, ELOF1 is already an intrinsic part of the elongating complex in unperturbed conditions where it stimulates transcription elongation (Fig. 2). Its dual function as an elongation and repair factor can be the cause of the embryonic lethality observed in ELOF1 KO mice ^61^ and may explain why thus far no ELOF1 mutations were found in TC-NER related syndromes, something commonly observed for other TC-NER factors ^12^. The role of ELOF1 in TC-NER is highly conserved, also in yeast in which TC-NER is differently organized. For example, in yeast TBLs can be repaired in a Rad26-independent manner ^50^, and no homolog of UVSSA is detected. As ELOF1 is an integral part of the elongation complex we speculate that this is the reason why ELOF1 is crucial in both Rad26-dependent and -independent repair pathways, since its presence in the stalled Pol II complex is not dependent on Rad26.

ELOF1 promoted UVSSA binding to lesion-stalled Pol II, resulting in subsequent TFIIH recruitment, which promotes assembly of the full incision complex to excise the TBL and restart transcription. In the absence of ELOF1, TC-NER can still be initiated since CSB and the CRL4^CSA^ E3 ubiquitin ligase complex are still properly recruited to lesion-stalled Pol II (Fig 5H). Interestingly, although UVSSA was previously shown to be incorporated into the TC-NER complex through a direct interaction with CSA ^56,62^, we found that in the absence of ELOF1, a repair intermediate accumulated that consists of CSA but not of UVSSA. This suggests that more control steps are needed to recruit or stably incorporate UVSSA, and that this is not only mediated via a direct interaction with CSA, which is in line with the previously observed CSA-independent UVSSA recruitment ^57,63^. Such tight regulation will control the subsequent TFIIH recruitment and assembly of the incision complex and may represent an important proof-reading step that prevents build-up of the incision complex on non-lesion stalled Pol II. An example of such a regulatory mechanism is the recently discovered TBL-induced ubiquitylation of a single lysine residue (K1268) of Pol II, that is crucial for Pol II stability and TFIIH recruitment ^18,19^. Interestingly, based on recent structural analysis of the elongation complex in yeast ^32^, the K1268 ubiquitylation site is in close proximity of ELOF1 (Suppl. Fig. 10). In the absence of ELOF1, Pol II ubiquitylation is reduced. We hypothesize that ELOF1 might either stimulate this ubiquitylation by facilitating a correct orientation of the elongation complex, or is involved in recruiting E3 ligases or repair factors that promote Pol II ubiquitylation. As UVSSA was shown to be important for the K1268 ubiquitylation ^18^ and its recruitment is promoted by ELOF1, this might argue for the latter.

Our data, together with recent cryo-EM studies, indicate that TC-NER factors embrace the complete elongation complex, with CSB binding to upstream DNA extruding from Pol II ^13^ and ELOF1 binding to downstream DNA entering Pol II ^32^ (Suppl. Fig. 10). As CSB promotes the forward translocation of Pol II, thereby sensing for a TBL ^13^, it is tempting to speculate that the presence of ELOF1 at the opposite site of Pol II might promote Pol II backtracking upon DNA damage, thereby facilitating repair factors to access the TBL. Since TFIIH might be involved in this backtracking process ^1^, this could explain why TFIIH recruitment is reduced upon ELOF1 depletion.

In addition to its role in TC-NER, our data show that ELOF1 plays an important role in preserving genome stability upon DNA damage, likely by preventing transcription-mediated replication stress (Fig. 5). The chromatin binding of the transcription machinery and the inability to clear Pol II from the DNA is assumed to play an important role in the onset of transcription-replication conflicts ^8,61,64^. Even though CSB and ELOF1 depletion had similar effects on the prolonged binding of Pol II upon DNA damage (Fig. 4A), only ELOF1 KO resulted in a clear replication defect and increased genome instability (Fig. 5E,F). This suggests that Pol II is differently processed in the absence of ELOF1 compared to what happens in CSB-deficient cells. This is most likely not caused by Pol II degradation, as loss of ubiquitylation is observed in the absence of both ELOF1 and CSB (Fig. 4I and Suppl. Fig. 8J). This implies that ELOF1 either has a function in Pol II release upon stalling at a lesion or that, in the absence of ELOF1, Pol II cannot be properly released from the DNA by incoming replication forks, resulting in an increase in transcription-replication conflicts (Fig. 5H). Together, our results show that ELOF1 is an important guardian of elongating Pol II by protecting transcription from the severe consequences of TBLs via two mechanisms; by stimulating repair and by preventing transcription-replication conflicts.

## Acknowledgments

We thank the Optical Imaging Centre and the proteomics center of the Erasmus Medical Center for support with microscopes and mass spectrometry analysis. We thank the Advanced Sequencing Facility of the Francis Crick Institute’s for technical assistance on the DRB/TT_chem_-seq.

## Author contributions

M.E.G performed the majority of the experiments and generated ELOF1 and CSB KO cell lines and ELOF1- and RPB1-KI cell lines. D.Z. generated CSB- and UVSSA-KI cells and performed live-cell imaging experiments and the TCR-UDS. K.S., D.A.P., W.G, S.L, and J.J.W. performed and supervised all experiments in *S.cerevisiae*. B.S. and M.E.G. performed the CRISPR/cas9 screen, and B.E. and R.B. analyzed and supervised the screen. C.M. and A.R.C. performed and supervised the metaphase spread and DNA fiber analysis. S.C., R.M., and J.Q.S. performed and supervised the DRB/TT_chem_-seq. M.v.T. performed the alamar blue cell viability assay. M.v.d.W. and H.L. performed and supervised the experiments in *C. elegans*. R.J. provided experimental support. J.L. generated images of Pol II structure. B.G. performed Monte-Carlo based-modeling and was supervised by A.H.. K.B. and J.A.A.D. performed and supervised mass spectrometry analysis. A.R. performed UDS experiments and A.F.T. performed FACS sorting, both supervised by W.V.. J.A.M. conceived and supervised the project and together with M.E.G. wrote the manuscript with input from all authors.

## Data and materials availability

All DRB/TT_chem_-seq data used in this study is available under GEO accession: GSE148844. All CPD-seq data is available under GEO accession: GSE149082. Raw mass spectrometry data is available upon request.

## Methods

### Cell lines and cell culture

MRC-5 (SV40) immortalized human lung fibroblast cells and HCT116 colorectal cancer cells were cultured in a 1:1 mixture of DMEM (Gibco) and Ham’s F10 (Invitrogen) supplemented with 10% fetal calf serum (FCS, Biowest) and 1% penicillin-streptomycin in a humidified incubator at 37°C and 5% CO_2_. C5RO fibroblasts (hTert), CS3BE (CS-A, SV40), XP186LV (XPC-/-) and CS216LV (CS-A, hTert) cells were maintained in Ham’s F10 with 15% FCS and antibiotics.

For stable isotope labeling of amino acids in culture (SILAC), cells were grown for two weeks (>10 cell doublings) in arginine/lysine-free SILAC DMEM (Thermofisher) supplemented with 15% dialyzed FCS (Gibco), 1% penicillin-streptomycin, 200 µg/ml proline (Sigma), and either 73 μg/mL light [^12^C_6_]-lysine and 42 μg/mL [^12^C_6_, ^14^N_4_]-arginine (Sigma) or heavy [^13^C_6_]-lysine and [^13^C_6_, ^15^N_4_]-arginine (Cambridge Isotope Laboratories).

HCT116 knock-out cells were generated by transiently transfecting HCT116 cells with a pLentiCRISPR.v2 plasmid ^23^ containing appropriate sgRNAs. Transfected cells were selected using 1 µg/ml puromycin (Invitrogen) for 2 days and single cells were seeded to allow expansion. Genotyping of single-cell clones was performed by immunoblotting or genomic PCR as indicated. sgRNAs sequences can be found in table 1, see below.

**Table 1.**
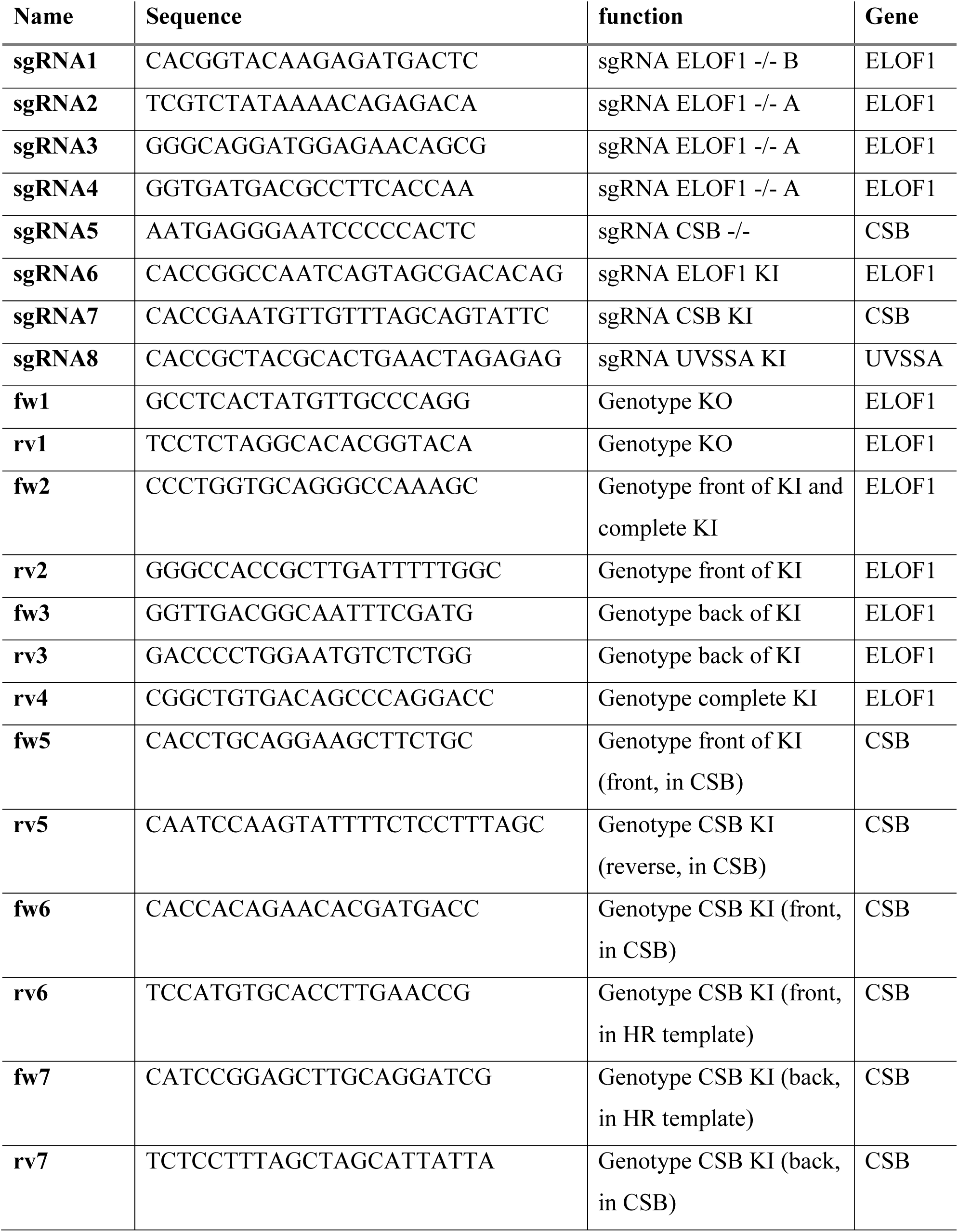
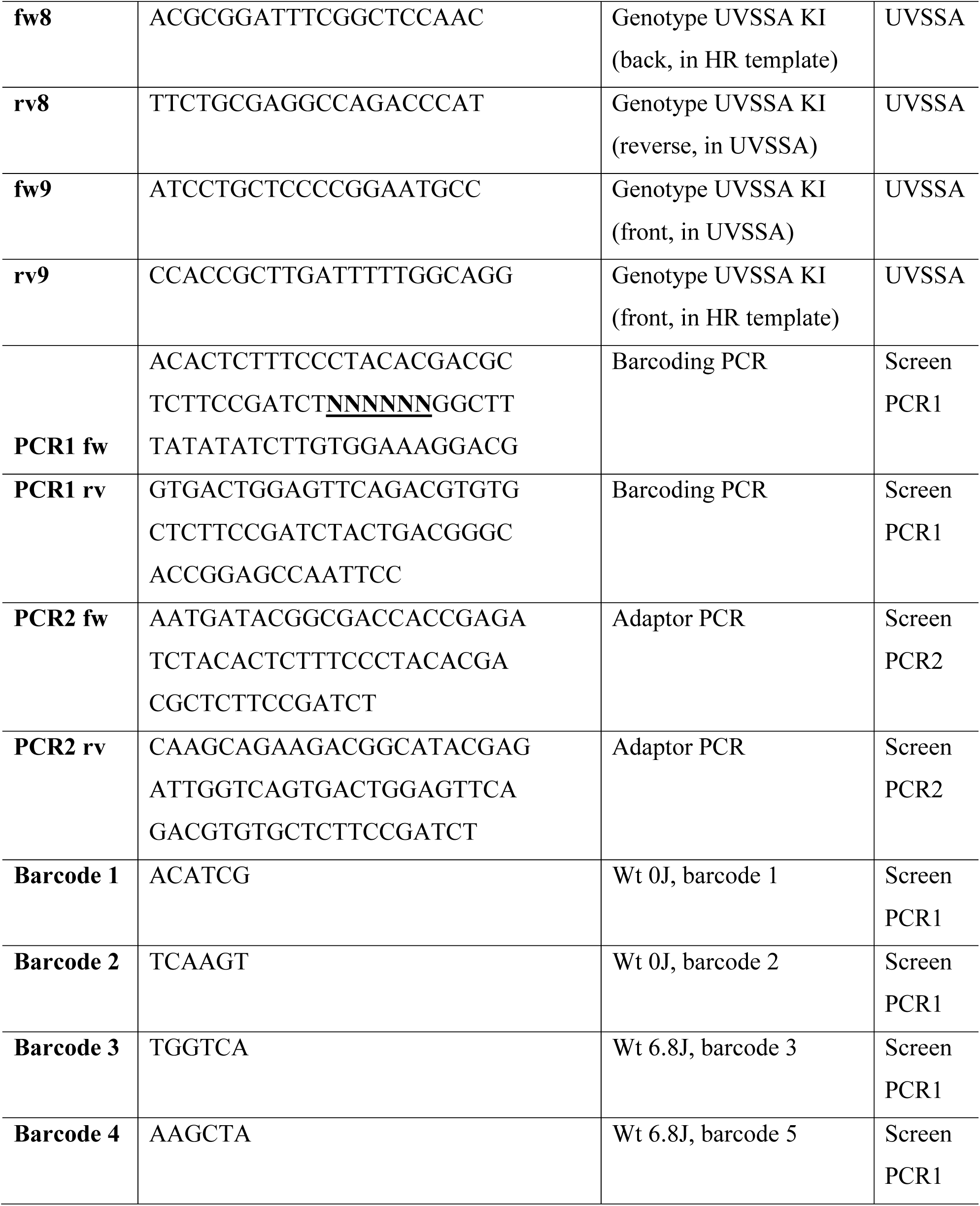
Primers.

ELOF1 complemented cell lines were generated by lentiviral transduction in ELOF1 -/- cells. Therefore, full-length expression constructs with ELOF1-Flag-GFP, or Wt or mutated ELOF1-Flag were synthesized (Genscript) and inserted in a pLenti-CMV-puro-DEST plasmid ^65^. After transduction, cells were selected with 1 µg/ml puromycin.

HCT116 osTIR1 knock-in (KI) cells ^66^ were generated by transiently transfecting cells with an sgRNA-containing pLentiCRISPR.v2 plasmid (sgRNA sequences in table 1, see below) targeting the stop codon of *ELOF1, CSB* or *UVSSA* and co-transfecting a homology-directed repair template, which included an Auxin-inducible Degron, fluorescent mScarletI-tag, HA-tag, hygromycin resistance cassette and homology arms (140 bp for ELOF1, 200 bp for CSB and UVSSA, sequence upon request) ^67^. Subsequently, cells were seeded in a low density to allow expansion and were kept in presence of 100 µg/ml hygromycin for two weeks to select for successful recombination. Single-cell clones were genotyped and homozygous KI clones were selected for further analysis. A GFP-RPB1 KI was generated in HCT116 Wt or ELOF1-KI cells as previously described by Steurer *et al.* ^68^. MRC-5 GFP-RPB1 KI cells ^68^ expressing CPD-PL-mCherry were generated as described previously ^69^.

Genotyping PCR was performed on genomic DNA (isolated using a PureLink™ Genomic DNA Mini Kit according to manufacturer’s protocol) with Phusion (NEB) or taq (Invitrogen) polymerases according to manufacturer’s protocol. Primer sequences can be found in table 1, see below. If necessary for assessing genomic alterations, PCR fragments were sequenced with forward primers and indels were analyzed using TIDE analysis ^70^.

siRNA transfections were performed 2 or 3 days before each experiment using Lipofectamine RNAiMax (Invitrogen) according to manufacturer’s protocol. siRNAs were purchased from Dharmacon: siELOF1 #1: 5’-CCGUGUGCCUAGAGGAAUUUU-3’, siELOF1 #2: 5’-GAAAUCCUGUGAUGUGAAAUU-3’, siCSB: 5’-GCAUGUGUCUUACGAGAUAUU-3’, siXPF: M-019946-00, siSPT4: L-012602-00-0005, siSPT5: L-016234-00-0005, siCSA: L-011008-00-0005. Knock-down efficiency was determined by immunoblot or RT-qPCR.

For UV-C irradiation, cells were washed with PBS, and placed under a 254 nm germicidal UV-C lamp (Philips). Duration of irradiation was controlled with an air-pressured shutter connected to a timer and cells were irradiated with doses as indicated. Cells were treated with VCP inhibitor (Seleck Chemicals, 5 µM) directly after UV irradiation or pre-treated 1 hour before irradiation with proteasome inhibitor MG132 (Enzo, 50 µM) or NEDD8 E1 Activating Enzyme Inhibitor (NAEi) MLN4924 (R&D systems, 10 µM) where indicated. Cell were treated for 1 hour with the following chemicals: Actinomycin D (Sigma, 1 µg/ml), Flavopiridol (Sigma, 1 µM), THZ1 (Xcessbio, 2 µM), Mitomycin C (Sigma, 10 µg/ml unless indicated differently), or potassium bromate (KBrO_3_, Sigma). Cells were exposed continuously to camptothecin or treated for 24 hours with cisplatin, illudin S, or hydroxyurea (all Sigma). Final concentrations of all inhibitors were diluted in culture media and cells were washed once with PBS before putting fresh media after removing damaging agent when necessary. For ionizing radiation, plates were irradiated using an RS320 X-ray cabinet (X-Strahl). For photoreactivation, cells were washed with PBS and covered with a thin layer of HBSS (Thermofisher) before exposing them to white-light tubes (General Electric Lighting Polylux LX F36W/840) for 10 minutes at 37 °C ^69^. Mock-treated samples were covered with tinfoil during photo-reactivation.

### GeCKO v2 lentiviral library production and transduction

We used the lentiCRISPRv2 human library designed by Shalem *et al.* ^71^ and obtained from Addgene. The sgRNA library was synthesized using array synthesis as previously described ^71^ and cloned as a pool into the lentiCRISPR transfer plasmid for virus production.

To produce the pooled lentiviral library, twelve T-225 flasks of HEK293T cells were seeded at ∼40% confluency the day before transfection. Per flask 10 µg of pVSVg, and 15 µg of psPAX2 (Addgene) packaging plasmids and 20 µg of lentiCRISPR plasmid library were transfected using Lipofectamine 2000 and Plus reagent (Life Technologies), according to manufacturer’s instructions. After 6 hours the medium was changed, and after 60 hours the medium was collected and centrifuged at 3,000 rpm for 10 minutes at 4 °C to pellet cell debris. The supernatant was filtered through a 0.45 µm low protein-binding membrane (Millipore Steriflip HV/PVDF). To achieve a 300 times concentration of the GeCKO pooled library, the virus was ultracentrifuged (Sorvall) at 24,000 rpm for 2 hours at 4 °C and then resuspended overnight at 4 °C in D10 supplemented with 1% BSA. Aliquots were stored at –80°C.

Per condition 20 million MRC-5 cells were transduced at 75% confluency in 145 cm^2^ dishes with concentrated lentivirus diluted in 18 ml of culture medium supplemented with 12 µg/mL polybrene (Sigma). The virus titer was determined to achieve a multiplicity of infection of <0.25. The next day, cells were re-seeded at 25% confluency in culture medium containing 2 µg/ml Puromycin. Cells were expanded for 1 week in puromycin-containing medium. Culture medium was refreshed every other day.

### Genome-wide CRISPR screen

For UV irradiation or mock treatment 30 million transduced and puromycin-selected cells were seeded per condition at 40% confluency in 145 cm^2^ dishes (2.25 million cells per dish) in medium without puromycin. The next day (day 0) dishes were mock-treated or irradiated with 6.8 J/m^2^ UV-C. Control cells (mock-treated) and UV irradiated cells were washed with PBS and (mock) irradiated every day for 10 consecutive days. The culture medium was refreshed after each irradiation. Mock-treated cells were reseeded to 40% confluency when they reached a confluency > 90%. After the last irradiation cells were given 24 hours to recover and gDNA was isolated using the Blood & Cell Culture DNA Midi Kit (Qiagen) according the manufacturers protocol (DNA content of MRC-5 cells was estimated at 10 pg per cell, genomic DNA of max 15 million cells was loaded per column). The screen was performed in duplicate.

### PCR and next-generation sequencing

Per condition, sgRNA sequences of at least 300 µg of DNA (of ∼30 million cells) were amplified by PCR (PCR1) using barcoded forward primers to be able to deconvolute multiplexed samples after next-generation sequencing (primers and barcodes are listed in table 1, see below). PCR1 was performed on 3 µg of gDNA in a total volume of 50 µl per reaction. Each PCR1 reaction contained 1 U of Phusion Hot Start II Polymerase (Thermo Fisher Scientific), 1x reaction buffer, 200 nm of each dNTP, 0.5 µM of both forward and reverse primer, and 3% DMSO. The following PCR program was used: initial denaturation for 3 minutes at 98°C; 35 cycles of denaturation for 1 sec at 98°C, primer annealing for 30 sec at 60°C, extension for 30 sec at 72°C, and final extension of 10 minutes at 72°C. Individual PCR reaction products were pooled per condition and 2 µl of pooled PCR product was used for a second PCR (PCR2) using primers containing adapters for next-generation sequencing (table 1, see below). The same PCR program was used as for PCR1, except that only 15 cycles were applied. 30 µl of PCR2 product was cleaned up to remove primer pairs using the NucleoSpin Gel & PCR clean up kit (Bioké). Equal DNA content between conditions was checked by gel electrophoresis and samples were equimolarly pooled and subjected to Illumina next-generation sequencing as described before ^24^. Mapped read-counts were subsequently used as input for the Model-based Analysis of Genome-wide CRISPR-Cas9 Knockout (MAGeCK) analysis software package, using version 0.5. For each condition, two biological replicates were performed. All conditions were sequenced simultaneously. To determine which genes showed a significant negative selection after 10 days of UV treatment, the sequencing data were analyzed with the MAGeCK tool ^25^. Gene ontology (GO) term enrichment analysis was performed using the g:Profiler website. Genes with a FDR<0.1 were analyzed and the top 10 biological processes affected by UV were identified.

### Survival assays

For clonogenic survival assay, 200-300 cells were seeded per well in triplicate in a 6-well plate. The following day, cells were treated with different DNA damaging agents. Following treatment, colonies were grown for 7 to 10 days after which they were fixed and stained using Coomassie blue (50% methanol, 7% acetic acid and 0.1% coomassie blue (all Sigma)). To assess the growth speed of siRNA-transfected cells, 10,000 (HCT116) or 20,000 (ELOF1 -/-A) cells were seeded in a 6-well plate and grown for 10 days after transfection. Colony numbers were counted using GelCount (Oxford Optronix Ltd.). Relative colony number was plotted of at least 2 independent experiments, each performed in triplicate. Levels were normalized to mock-treated, set to 100 and plotted with SEM. Statistics was performed using independent T-test.

For AlamarBlue survival assay, siRNA-transfected cells were seeded to confluency in presence of 0.5% serum in triplicate in 96-well plates to arrest cells in G_0,_ and UV-irradiated after 30 hours. 72 hours after UV irradiation, AlamarBlue® (Invitrogen) was added for 4 hours and fluorescence was measured at 570 nm using a SpectraMax iD3 reader. Data were background corrected and normalized to mock-treated conditions.

### RNA isolation, cDNA synthesis and RT-qPCR

To determine ELOF1 expression levels, RNA was isolated using the RNeasy mini kit (Qiagen) and cDNA was synthesized using the SuperScript™ II Reverse Transcriptase (Invitrogen), both according to the manufacturer’s protocol. The generated cDNA was amplified using 1x taqman assay (ELOF1: Hs00361088_g1, GAPDH: 4333764T, both Thermofisher) and 1x taqman gene expression master mix (Thermofisher) by activating UNG for 2 minutes at 50°C, activating the polymerase for 10 minute at 95°C, followed by 40 cycles of 15 seconds of denaturing at 95°C and 1 minute of annealing and extending at 60°C in a CFX96 Touch Real-Time PCR Detection System. mRNA expression levels were normalized to GAPDH using the 2^-ΔΔCt^ method^72^.

### Cell lysis and immunoblotting

Cells were directly lysed in SDS Page loading buffer (0.125M Tris pH 6.8, 2% SDS, 0.005% bromophenol blue, 21% glycerol, 4% β-mercaptoethanol) or, for assessing the chromatin fraction, one confluent 9.6 cm^2^ dish was lysed for 30 minutes at 4°C in buffer containing 30 mM HEPES pH 7.5, 130 mM NaCl, 1 mM MgCl_2_, 0.5% Triton X-100, cOmplete^™^ EDTA-free protease inhibitors (Roche), Phosphatase inhibitor cocktail 2 (Sigma), N-ethylmaleimide (Sigma), and 50 µM MG132i. Chromatin was pelleted at 15,000 for 10 minutes at 4°C and washed once. Finally, the chromatin was digested for 30 minutes at 4°C in presence of 50 U of benzonase (Millipore) before adding SDS Page loading buffer and incubating 5 minutes at 95°C. Chromatin fractions or cell lysates were separated on 4-15% Mini-PROTEAN TGX^TM^ Precast Protein Gels (BioRad). Proteins were transferred onto PVDF membranes (0.45µm, Merck Millipore) at 4°C, either 1.5h at 90V with 1x transfer buffer (25mM TRIS, 190mM Glycine, 10% methanol) or overnight at 25V in 2x transfer buffer (50mM TRIS, 380mM Glycine). Membranes were blocked with 5% BSA (Sigma) in PBS-tween (0.05%) and probed with primary antibodies (Table 2, see below). Subsequently, membranes were extensively washed with PBS-tween and incubated with secondary antibodies coupled to IRDyes (LI-COR, table 3, see below) to visualize proteins using an Odyssey CLx infrared scanner (LI-COR).

**Table 2:** primary antibodies.

| **Antibody** | **Host** | **Source** | **Dilutions** |  |
| --- | --- | --- | --- | --- |
|  |  |  | **WB** | **IF** |
| 53BP1 | Rb | Santa Cruz, sc-22760 | N.A. | 1/1000 |
| BrdU (CldU) | Rat | Abcam, ab6326 | N.A. | 1/500 |
| BrdU (IdU) | Ms | BD Biosciences, B44, 347580 | N.A. | 1/100 |
| CSA/ERCC8 | Ms | Santa Cruz, sc376981 | 1/250 | N.A. |
| CSB/ERCC6 | G | Santa Cruz, sc10459 | 1/500 | N.A. |
| CSB/ERCC6 | Rb | Antibodies-online, ABIN2855858 | 1/1000 | N.A. |
| GFP | Ms | Roche, 14314500 | 1/1000 | N.A. |
| GFP | Rb | Abcam, Ab290 | 1/1000 | N.A. |
| HA | R | Roche, 11867423001 | 1/1000 | N.A. |
| Lamin B1 | Rb | Abcam, | 1/1000 | N.A. |
| p62/GTF2H1 | Ms | Sigma Aldrich, WH0002965M1 | 1/1000 | N.A. |
| RPB1 (Pol II) | Rb | Cell signalling, D8L4Y | 1/1000 | N.A. |
| RPB3 | Rb | Abcam, ab138436 | 1/1000 | N.A. |
| RPB9 | Rb | Abcam, ab192407 | 1/500 | N.A. |
| Ser2 | R | Chromotek, 3E10 | 1/1000 | N.A. |
| SPT4/SUPT4H1 | Rb | Cell signalling, D3P2W | 1/1000 | N.A. |
| SPT5/SUPT5H | Rb | Bethyl, A300-869A | 1/500 | N.A. |
| SSRP1 | Ms | Biolegend, 609701 | 1/1000 | N.A. |
| Tubulin | Ms | Sigma Aldrich, B512 | 1/5000 | N.A. |
| XPD | Ms | Abcam, ab54676 | 1/1000 | N.A. |
| XPF | Ms | Santa Cruz, sc-136153 | 1/500 | N.A. |

**Table 3:** secondary antibodies.

|  |  |  | **Dilutions** |  |
| --- | --- | --- | --- | --- |
| **Antibody** | **Host** | **Source** | **WB** | **IRDye** |
| Rabbit | goat | Sigma, sab4600215 | 1/10000 | 770 |
| Rabbit | goat | Sigma, sab4600200 | 1/10000 | 680 |
| Mouse | goat | Sigma, sab4600199 | 1/10000 | 680 |
| Mouse | goat | Sigma, sab4600214 | 1/10000 | 770 |
| Goat | donkey | Sigma, sab4600375 | 1/10000 | 770 |
| Rat | goat | Sigma, sab4600479 | 1/10000 | 770 |
| **Antibody** | **Host** | **Source** | **IF** | **AlexaFluor** |
| Rabbit | donkey | Invitrogen, A21207 | 1/1000 | 594 |
| Mouse | goat | Invitrogen, A11001 | 1/300 | 488 |
| Rat | donkey | Jackson Immuno-Research<br>lab., 712-166-153 | 1/150 | Cy3 |

### Fluorescence Recovery After Photobleaching (FRAP)

For FRAP, a Leica TCS SP5 microscope (LAS AF software, Leica) equipped with a HCX PL APO CS 63x 1.40 NA oil immersion lens (ELOF1, RPB1, CSB) or Leica TCS SP8 microscope (LAS AF software, Leica) equipped with a HC PL APO CS2 63x 1.40 NA oil immersion lens (UVSSA) was used. Cells were maintained at 37°C and at 5% CO_2_ during imaging. A narrow strip of 512 x 32 pixels (for ELOF1 and RPB1) or 512×16 (for CSB and UVSSA) spanning the nucleus was imaged every 400 ms (200 ms for UVSSA during pre-bleach) at 400 Hz using a 488 nm laser (RPB1) or 561 nm laser (ELOF1, CSB, UVSSA). 25 (RPB1), 40 (ELOF1), or 5 (CSB, UVSSA) frames were measured to reach steady state levels before photobleaching (1 frame 100% laser power for RPB1 and ELOF1, 2 frames for CSB and UVSSA). After photobleaching, the recovery of fluorescence was measured with 600 (ELOF1 and RPB1), 40 (CSB) or 20 (UVSSA) frames until steady-state was reached. Fluorescence intensity was measured inside and outside of the nucleus and recovery was determined by correcting for background signal and normalizing the values to the average pre-bleach fluorescence intensities. Relative fluorescence intensity levels were calculated using the pre-bleach intensities corrected for background. Immobile fractions (F_imm_) were calculated using the individual and average (indicated by <brackets>) fluorescence intensities after bleaching (I_bleach_) and fluorescence intensities after recovery from the bleaching (I_recovery_):

*F_imm_*= 1 −(*I_recovery,UV_* − < *I_bleach_* >)/(< *I_recovery,unc_* > − < *I_bleach_* >)

Experimental FRAP curves of Pol II were simulated using Monte-Carlo-based computational modeling as described previously ^68^ to determine the residence time of elongating Pol II and the fraction size of promoter-bound and elongating Pol II.

### Native immunoprecipitation (IP)

Cells were mock-treated or irradiated with 16 J/m^2^ UV-C 1 hour prior to cell harvest. Cell pellets were prepared from 3 confluent 145 cm^2^ dishes per condition for IP followed by immunoblot or 8 confluent 145 cm^2^ dishes per condition for mass spectrometry. Cells were collected by trypsinization and pelleted in cold PBS using centrifugation for 5 minutes at 1500 rpm. After one wash with cold PBS, cell pellets were stored at −80°C until immunoprecipitation.

For immunoprecipitation, pellets were thawed on ice and lysed for 20 minutes at 4°C in HEPES buffer containing 30 mM HEPES pH 7.6, 1 mM MgCl_2_, 150 mM NaCl, 0.5% NP-40, and 1x cOmplete^™^ EDTA-free Protease Inhibitor Cocktail (Roche). Chromatin was pelleted by spinning 5 minutes at 10,000 g at 4°C and subsequently incubated for 1 hour at 4°C in HEPES buffer containing 500 units of Benzonase (Millipore) and 2 µg Pol II antibody (ab5095, abcam) or IgG (sc2027, Santacruz) to digest the chromatin. After 1 hour, the NaCl was increased to 300 mM to inactivate benzonase and antibody-binding was continued for another 30 minutes. The undigested fraction was pelleted at 13,200 rpm for 10 minutes at 4°C and the soluble, antibody-bound fraction was immunoprecipitated for 90 minutes at 4°C using 25 µL slurry salmon sperm protein A agarose beads (Millipore). Unbound proteins were removed by washing the beads 5 times in wash buffer (30 mM HEPES pH 7.6, 150 mM NaCl, 1mM EDTA, 0.5% NP-40, and 0.2x cOmplete^™^ EDTA-free Protease Inhibitor Cocktail). Bound proteins were eluted in SDS page loading buffer and separated on 4-15% Mini-PROTEAN TGX^TM^ Precast Protein Gels (BioRad). Samples were processed for immunoblotting or fixed and stained for mass spectrometry using Imperial protein stain (Pierce) according to manufacturer’s protocol.

For ELOF1 IP, the same protocol was followed but instead of adding antibody during chromatin digestion, precipitation was performed using RFP-Trap® agarose beads (Chromotek) and binding control agarose beads (Chromotek).

### Cross-linked immunoprecipitation

Cells were mock-treated or irradiated with 16 J/m^2^ UV-C one hour prior to cell harvest. Cell pellets were prepared from 8 confluent 145 cm^2^ dishes per condition for mass spectrometry. MRC-5 GFP-RPB1 KI cells were used for Pol II IP (Flag-beads) and HCT116 ELOF1-KI cells were used for ELOF1 IP (HA-beads).

Crosslinked IP was performed as described previously ^73^ with modifications as indicated. Cells were cross-linked with 1% paraformaldehyde (PFA) in serum-free DMEM for 7 minutes with constant shaking before quenching the reaction for 5 minutes with glycine (final concentration of 0.125 M). Cells were collected by scraping in PBS with 10% glycerol and 1 mM PMSF and pelleted for 15 minutes at maximum speed at 4°C. Consequently, chromatin was purified by washing the cell pellets for 30 minutes at 4°C in buffer 1 (50 mM HEPES, 150 mM NaCl, 1 mM EDTA, 0.5 mM EGTA, 0.25% Trition X-100, 0.5% NP-40, 10% glycerol), pelleting the cells 10 minutes at 1300 rpm, washing the pellet twice with buffer 2 (10 mM Tris pH 8.0, 200 mM NaCl, 1 mM EDTA, 0.5 mM EGTA) and finally pelleting the chromatin, all at 4°C. Chromatin was sonicated in RIPA buffer (10 mM Tris pH 7.5, 150 mM NaCl, 5 mM EDTA, 0.1% SDS, 1% Sodium Deoxycholate and 0.5 mM EGTA) using the Bioruptor Sonicator (Diagenode) with 14 cycles of 15s on/15s off using the highest amplitude. Extracted chromatin was collected by spinning 15 minutes at maximum speed and pre-cleared for 30 minutes with Protein G agarose beads (Pierce) at 4°C. IP was performed by incubating 4 hours at 4°C with Flag M2 agarose beads (Sigma). Finally, aspecific interactors were removed by washing five times with RIPA buffer and proteins were eluted and crosslinking was reversed by incubating 30 minutes at 95°C in SDS Page loading buffer. Samples were separated on 4-15% Mini-PROTEAN TGX^TM^ Precast Protein Gels (BioRad) and fixed and stained using imperial protein stain in preparation of mass spectrometry. To all buffers, 1 mM PMSF, 0.5 mM Na_2_VO_4_, 5 mM NaF, 5 mM NaPPi, 10 mM β-glycerol and cOmplete^™^ EDTA-free Protease Inhibitor Cocktail were added.

For ELOF1 IP, the same protocol was followed with minor alterations. Cells were crosslinked in 1 mM dithiobis(succinimidyl propionate) (DSP) in PBS for 30 minutes and quenched by adding Tris pH 7.5 to a final concentration of 25 mM for 10 minutes. IP was performed using HA-agarose beads (Sigma) and beads were incubated for 5 minutes at 95°C to elute and reverse cross-linked immunocomplexes.

### Mass spectrometry

SDS-PAGE gel lanes were cut into slices and subjected to in-gel reduction with dithiothreitol (Sigma, D8255), alkylation with iodoacetamide (Sigma, I6125) and digestion with trypsin (sequencing grade; Promega) as previously described ^57^. Nanoflow liquid chromatography tandem mass spectrometry (nLC-MS/MS) was performed on an EASY-nLC 1200 coupled to a Lumos Tribid Orbitrap mass spectrometer (ThermoFisher Scientific) operating in positive mode. Peptide mixtures were trapped on a 2 cm x 100 μm Pepmap C18 column (Thermo Fisher 164564) and then separated on an in-house packed 50 cm x 75 μm capillary column with 1.9 μm Reprosil-Pur C18 beads (Dr. Maisch) at a flowrate of 250 nL/min, using a linear gradient of 0–32% acetonitrile (in 0.1% formic acid) during 90 min. The eluate was directly sprayed into the electrospray ionization (ESI) source of the mass spectrometer. Spectra were acquired in continuum mode; fragmentation of the peptides was performed in data-dependent mode by HCD. Mass spectrometry data were analyzed using the MaxQuant software (version 1.6.3.3). The false discovery rate (FDR) of both PSM and protein was set to 0.01 and the minimum ratio count was set to 1. The Andromeda search engine was used to search the MS/MS spectra against the UniProt database (taxonomy: Homo sapiens, release June 2017), concatenated with the reversed versions of all sequences. A maximum of two missed cleavages was allowed. In case the identified peptides of two proteins were the same or the identified peptides of one protein included all peptides of another protein, these proteins were combined by MaxQuant and reported as one protein group. Before further analysis, known contaminants and reverse hits were removed. Gene ontology (GO) term enrichment analysis was performed using the g:Profiler website. Genes with an average SILAC ratio of >2.5 were analyzed and the top 10 biological processes affected by UV were identified.

### DRB/TT_chem_-seq method

The DRB/TT_chem_-seq was carried out as described in Gregersen *et al.* ^40^ in two biological replicates. Briefly, 8 × 10^6^ cells were incubated in 100 µM DRB (Sigma-Aldrich) for 3.5 hours. The cells were then washed twice in PBS and fresh, DRB-free medium was added to restart transcription. The RNA was labelled *in vivo* with 1 mM 4SU (Glentham Life Sciences) for 10 minutes prior to the addition of TRIzol (Thermo Fisher Scientific), which was used to stop the reaction at the desired time point. Following extraction, 100 µg of RNA was spiked-in with 1 µg 4-thiouracile labelled S. cerevisiae RNA (strain BY4741, MATa, his3D1, leu2D0, met15D0, ura3D0), and then fragmented with NaOH and biotinylated with MTSEA biotin-XXlinker (Biotium). The biotinylated RNA was then purified using µMACS Streptavidine MicroBeads (Miltenyi Biotec) and used for library preparation. The libraries were amplified using the KAPA RNA HyperPrep kit (Roche) with modifications as described in ^74^.The fragmentation step was omitted and the RNA, resuspended in FPE Buffer, was denatured at 65°C for 5 min. Two SPRI bead purifications were carried out, with a bead-to-sample volume ratio of 0.95x and 1x, respectively. The libraries were then sequenced with single end 75bp reads on the Hiseq4000, with ∼50,000,000 reads per sample.

### Computational Analysis

DRB/TT_chem_-seq data were processed using previously published protocol ^40^. Briefly, reads were aligned to human GRCh38 Ensembl 86. Read depth coverage was normalized to account for differences between samples using a scale factor derived from a yeast spike-in aligned and counted against Saccharomyces cerevisiae R64-1-1 Ensembl 86. ^75^Biological replicate alignments were combined for the purpose of visualization and wave-peak analysis in order to increase read-depth coverage.

A set of non-overlapping protein-coding genes (200kb+) were selected for wave-peak analysis. A meta-gene profile was calculated by taking a trimmed mean of each basepairs coverage in the region −2kb:+200kb around the TSS. This was further smoothened using a spline. Wave peaks were called at the maximum points on the spline, with the stipulation that the peak must advance with time before being subjected to manual review. Elongation rates (kb/min) were calculated by fitting a linear model to the wave peak positions as a function of time.

### EU incorporation

Cells were grown on coverslips and transcription levels were measured by pulse labeling with 5’ethynyl uridine (EU, Jena Bioscience) in Ham’s F10 medium supplemented with 10% dialyzed FCS and 20 mM HEPES buffer (both Gibco). Cells were labeled for 30 minutes using 400 µM EU (MRC-5 cells) or for 1 hour with 200 µM EU (HCT116 cells) before fixation with 3.7% formaldehyde (FA, Sigma) in PBS for 15 minutes at room temperature (RT). After permeabilisation with 0.1% Triton X-100 in PBS for 10 minutes and blocking in 1.5% BSA in PBS for 10 minutes, Click-it chemistry-based azide coupling was performed by incubation for 1 hour with 60 µM Atto594 Azide (Attotec, Germany) in 50 mM Tris buffer (pH 8) with 4 mM CuSO_4_ (Sigma), and 10 mM freshly prepared ascorbic acid (Sigma). DAPI (Brunschwieg Chemie) was added to visualize the nuclei. Coverslips were washed with 0.1% Triton in PBS and PBS only and mounted with Aqua-Poly/Mount (Polysciences). Cells were imaged with a Zeiss LSM 700 Axio Imager Z2 upright microscope equipped with a 40x Plan-apochromat 1.3 NA oil immersion lens or 63x Plan-apochromat 1.4 NA oil immersion lens (Carl Zeiss Micro Imaging Inc.). Integrated density of the EU signal in the nuclei was quantified using ImageJ. Therefore, the surface of each nucleus was determined based on the DAPI signal and mean fluorescence intensity was determined, corrected for the background signal. With these values, the integrated density was calculated, and plotted as single cell point with the average and SEM.

For assessing recovery of transcription after UV, cells were mock-treated or irradiated with 8 J/m^2^ UV-C 2 or 18 hours before EU incorporation. For recovery after mitomycin C, cells were mock-treated or incubated for 2 hours with 10 µg/ml Mitomycin C followed by a recovery period of 2 or 22 hours in normal medium. Integrated density was normalized to mock-treated.

### TC-NER-specific UDS

Amplified UDS was performed as described previously ^43^. Briefly, siRNA transfected primary XP186LV (XP-C patient cells) were serum-deprived for at least 24 hours in Ham’s F10 (Lonza) containing 0.5% FCS and antibiotics to arrest cells in G_0_. Cells were irradiated using 8 J/m^2^ UV and labelled for 7 hours with 20 µM 5-Ethynyl-2’-deoxyuridin (EdU) and 1 µM Floxouridine (Sigma). Subsequently, a 15-minute chase was performed with normal medium (0.5% FCS) supplemented with 10 µM thymidine (Sigma) to remove unincorporated EdU and cells were fixed and permeabilized with 3.7% FA and 0.5% Triton X-100 for 15 minutes. After permeabilizing the cells for 20 minutes with 0.5% Trition in PBS and washing with 3% BSA in PBS, endogenous peroxidase activity was quenched using 2% hydrogen peroxide (Sigma) for 15 minutes and incubated with PBS+ (0.5% BSA + 0.15 % glycine). Click-it chemistry was performed using the Click-it reaction cocktail containing Azide-PEG3-Biotin Conjugate (20 μM, Jena Bioscience), 1× Click-it reaction buffer (ThermoFisher Scientific), copper(III) sulfate (0.1 M) and 10× reaction buffer additive (ThermoFisher Scientific) for 1 hour and washed with PBS. To amplify the signal, coverslips were incubated for 1 hour using HRP-streptavidin conjugate (500 μg/ml), followed by PBS washes and a 10-minute incubation with Alexa-Fluor 488 labeled tyramide (100x stock, Thermofisher Scientific). Coverslips were washed with PBS and PBS+ and the nuclei were stained with DAPI in 0.1% triton. DAPI was washed away with 0.1% triton and slides were mounted using Aqua-Poly/Mount.

### Unscheduled DNA Synthesis (UDS)

Cells were grown to confluency on coverslips and serum-deprived (0.5%) for 2 days to arrest cells in G_0_. Cells were irradiated with 16 J/m^2^ and labeled with 20 µM EdU (Invitrogen) in Ham’s F10 supplemented with 10% dialyzed FCS and 20 mM HEPES buffer (both Gibco) for 3 hours before fixation for 15 minutes (3.7% FA and 0.5% triton X-100). Background signal was blocked by washing twice with 3% BSA in PBS for 10 minutes and nuclei were permeabilized for 20 minutes using 0.5% triton in PBS. EdU incorporation was visualized using Click-it chemistry, imaged and analyzed as described in the section *EU incorporation* with the adjustment that click-it reaction was performed for 30 minutes.

### Yeast strains

Yeast deletion strains used in this study are derivatives of the wild type strain BY4741 (*MATa his3Δ1 leu2Δ0 met15Δ0 ura3Δ0*) and Y452 (*MATα, ura3-52, his3–1, leu2-3, leu2-112, cir°*). The gene deletions were made by transformation of yeast cells with PCR products bracketing selection markers ^76^ or following published methods ^77^.

### Yeast UV sensitivity assay

Yeast cells were grown in YPD medium to mid-log phase. For spotting assay, cells were serially 10-fold diluted in fresh YPD medium and spotted on YPD plates. After exposure to different doses of UV-C light (254 nm), plates were incubated at 30^°^C in the dark and images were taken after 3-5 days of incubation. For quantitative UV survival assay, diluted yeast cells were plated on YPD plates and exposed to the indicated UV doses. The number of colonies on each plate was counted after incubating for 3 days at 30^°^C in the dark. The survival graph depicts the mean and SEM of three independent experiments.

### CPD-seq library preparation and sequencing

CPD-seq analysis of repair in Wt and *elf1*Δ mutant strains was performed as previously described ^49^. Briefly, yeast cells were grown to mid-log phase, pelleted, re-suspended in dH_2_O, and irradiated with 125 J/m^2^ UV-C light (254 nm). After UV treatment, cells were incubated in the dark in pre-warmed, fresh YPD medium for repair. Cells were collected before UV irradiation (No UV), immediately after UV (0 hours), and following a 2-hour repair incubation. The cells were pelleted and stored at −80^°^C until genomic DNA isolation.

Genomic DNA extraction, CPD-seq library preparation and quality control, sequencing with an Ion Proton sequencer, and data processing were performed as previously described ^49^. The resulting sequencing reads were aligned to the yeast genome (saccer3) using Bowtie 2 ^78^. Only CPD-seq reads associated with lesions at dipyrimidine sequences (i.e., TT, TC, CT, CC) were retained for further analysis.

Bin analysis for CPD repair along the transcribed strand (TS) and non-transcribed strand (NTS) of ∼4500 yeast genes was performed as previously described ^79^, using transcription start site (TSS) and polyadenylation site (PAS, also referred to as transcription termination site, TTS) coordinates from Park *et al.*^80^. A similar gene bin analysis was displayed for each yeast gene using the Java Treeview program ^81,82^. Genes were sorted by transcription rate ^83^. Single nucleotide resolution repair analysis adjacent to the TSS was performed as previously described ^79,84^. Nucleosome dyad coverage from MNase-seq experiments were obtained from Weiner *et al.*, ^85^ as reference. CPD-seq data for *elf1Δ* and Wt yeast was normalized using the fraction of CPDs remaining determined for bulk genomic DNA by T4 endonuclease V digestion and alkaline gel electrophoresis (see below).

### Analysis of bulk CPD repair in UV irradiated yeast

Alkaline gel electrophoresis to assay global DNA repair of bulk DNA was conducted as previously described ^86^. Yeast cell cultures were grown to mid-log phase in YPD media. Yeast cell cultures were briefly centrifuged to pellet, resuspended in dH_2_O, and exposed to 100 J/m^2^ UV-C light or left unirradiated for the “No UV” sample. Following irradiation, yeast cells were resuspended in YPD and incubated at 30°C. Aliquots were taken at each repair time point, briefly centrifuging to discard media supernatant prior to storing yeast cells at −80°C. Genomic DNA was isolated by bead beating the yeast cell pellets in 250 µL lysis buffer (2% Triton-X 100, 1% SDS, 100 mM NaCl, 10 mM Tris-Cl pH 8, 1 mM Na_2_EDTA) and 300 µL Phenol-Chloroform-Isoamyl alcohol (25:24:1). 300 µL TE pH 8 was added to each tube, briefly vortexing to mix. Samples were centrifuged and the DNA-containing aqueous layer was transferred to a fresh tube for ethanol precipitation. DNA pellets were resuspended in TE pH 8 containing 0.2 mg/mL RNase A, incubating at 37°C for 15 minutes prior to enzymatic digestion. Equal amounts of DNA were then treated with T4 endonuclease V (T4 PDG; NEB) and resolved by electrophoresis on a 1.2% alkaline agarose gel. Following neutralization and staining with SYBR Gold (Invitrogen), alkaline gels were imaged using the Typhoon FLA 7000 (GE Healthcare) and analyzed using ImageQuant TL 8.2 (GE Healthcare). The number of CPD lesions per kb was estimated using the ensemble average pixel density of each lane, corrected by the no enzyme control lane. Percent repair was calculated by normalizing the number of CPDs per kb to the no repair time point. Graphs represent the mean and SEM of at least 3 independent experiments.

### Repair analysis of UV induced CPDs in *RPB2* locus

Yeast cells were grown in synthetic dextrose (SD) medium at 30°C to late log phase (*A*_600_ ≈ 1.0), irradiated with 120 J*/*m^2^ of UV-C and incubated in YPD medium at 30°C in the dark. At different times of the repair incubation, aliquots were removed, and the genomic DNA was isolated. To map the induction and repair of UV-induced CPDs at the nucleotide resolution in a specific gene, libraries of DNA fragments adjoining the lesions were created by using the LAF-Seq (Lesion-Adjoining Fragment Sequencing) strategy ^87^ with some modifications. Briefly, the isolated genomic DNA was restricted with HincII and NruI to release a 553 bp *RPB2* gene fragment (168 bp upstream and 385 bp downstream of the transcription start site) and incised at the CPDs with T4 endonuclease V and treated *E. coli* endonuclease IV (New England Biolabs). The 3’ ends of the restricted and CPD-incised DNA fragments were ligated to Illumina sequencing adapters by using Circligase (Lucigen). After PCR amplification, the libraries were sequenced by using an Illumina HiSeq platform.

The sequencing reads were aligned to the *RPB2* gene by using Bowtie 2 ^78^. The numbers of reads from the UV-irradiated samples were normalized to those from the control (unirradiated) samples. Reads corresponding to CPDs at individual sites along the *RPB2* gene fragment were counted after subtraction of the background counts (in the unirradiated samples) by using codes in R. To more directly ‘visualize’ the CPD induction and repair profiles, images with band intensities corresponding to counts of aligned sequencing reads were created by using codes in R and MATLAB.

### *C. elegans* strains and UV sensitivity assays

*C. elegans* strains were cultured according to standard methods and outcrossed against Bristol N2, which was used as wild type. Mutant alleles were *xpc-1(tm3886)*, *csb-1(ok2335)*, and *elof-1(emc203)*. The loss of function *elof-1(emc203)* (Suppl. Fig. 6I) mutant strain was generated by injection of Cas9 protein together with tracrRNA and two crRNAs targeting *elof-1* (CAGTTGAATTGGGTGTCGAG and AGACGTCGATTGGCTCGGAG; Integrated DNA Technologies). Deletion animals were selected by genotyping PCR and sequencing. UV survival experiments were performed as described previously ^52^. Animals were irradiated at the indicated dose using two Philips TL-12 (40W) tubes emitting UV-B light. Briefly, ‘germ cell and embryo UV survival’ was determined by allowing UV-irradiated staged young adults to lay eggs on plates for 3 hours. To calculate the survival percentage, the total number of hatched and unhatched eggs was counted after 24 hours. For the ‘L1 larvae UV survival’, staged L1 larvae were UV irradiated and grown for 48 hours. Survival percentage was calculated by counting surviving animals that developed beyond the L2 stage and arrested animals as L1/L2 larvae.

### Metaphase spreads and chromosomal aberrations

Metaphase spreads were carried out as described previously ^88^. Briefly, cells were irradiated with 4 J/m^2^ or mock-treated 48 or 72 hours before preparing metaphase spreads (final confluence of 50-80%). Cells were arrested at metaphase by incubating with colcemid (*N*-methyl-*N*-deacetyl-colchicine, Roche, **10295892001**) for the last 14 hours before harvesting the cells. Collected cells were treated with hypotonic solution (KCl 0.075 M) for 30 minutes at 37 °C and fixed with methanol:acetic acid 3:1. Telomere-FISH was further carried out to study chromosomal aberrations. Metaphases were hybridized with telomere-repeat specific peptide nucleic acid (PNA) probes (Applied Biosystems) as described to label telomeres ^89^. A minimum 60 metaphase images were obtained using Carl Zeiss Axio Imager D2 microscope using 63x Plan Apo 1.4 NA oil immersion objective and analyzed with ImageJ software for chromosomal aberrations.

### DNA fiber analysis

DNA fiber analysis was carried out as described previously ^88,90^. Briefly, cells were sequentially pulse-labeled with 30 μM CldU (c6891, Sigma-Aldrich) and 250 μM IdU (I0050000, European Pharmacopoeia) for 15 min. For assessing fork progression after DNA damage, cells were irradiated with 4 J/m^2^ UV and incubated for 2 hours before pulse-labeling. After labeling, cells were collected and resuspended in PBS at 2.5 × 10^5^ cells per ml. The labeled cells were mixed 1:1 with unlabeled cells, and 2.5 µl of cells was added to 7.5 µl of lysis buffer (200 mM Tris-HCl, pH 7.5, 50 mM EDTA, and 0.5% (w/v) SDS) on a glass slide. After 8 min, the slides were tilted at 15–45°, and the resulting DNA spreads were air dried, fixed in 3:1 methanol/acetic acid overnight at 4 °C. The fibers were denatured with 2.5 M HCl for 1 hour, washed with PBS and blocked with 0.2% Tween-20 in 1% BSA/PBS for 40 min. The newly replicated CldU and IdU tracks were incubated (for 2.5 hours in the dark, at RT with anti-BrdU antibodies recognizing CldU and IdU (Table 2, see below), followed by a 1-hour incubation with secondary antibodies at RT in the dark: anti–mouse Alexa Fluor 488 and anti–rat Cy3 (Table 3, see below). Fibers were visualized and imaged by Carl Zeiss Axio Imager D2 microscope using 63X Plan Apo 1.4 NA oil immersion objective. Data analysis was carried out with ImageJ software. A one-way ANOVA was applied for statistical analysis using the GraphPad Prism Software.

### Immunofluorescence

Immunofluorescence was carried out as described previously ^91^. Cells were grown on 24-mm glass coverslips and mock-treated or irradiated with 8 J/m^2^ 48, 24 or 6 hours prior to fixation for 15 minutes in PBS with 3.7% FA. Subsequently, cells were permeabilized with 0.1% Triton X-100 in PBS and washed with PBS+ (0.15% BSA and 0.15% glycine in PBS). Cells were incubated for 2 hours at RT with rabbit anti-53BP1 antibody (table 2, see below) in PBS+. Thereafter, cells were washed with PBS+, 0.1% Triton and PBS+ before incubating 2 hours at RT with donkey anti-rabbit Alexa Fluor 594 conjugated antibody (table 3, see below) and DAPI. After washes with PBS+ and 0.1% Triton, coverslips were mounted with Aqua-Poly/Mount. Images were acquired with a Zeiss LSM700 Axio Imager Z2 upright microscope equipped with a 63x Plan-apochromat 1.4 NA oil immersion lens (Carl Zeiss Micro Imaging Inc.). Number of foci per nucleus was counted by using ImageJ.

---

**Supplemental figure 1.**
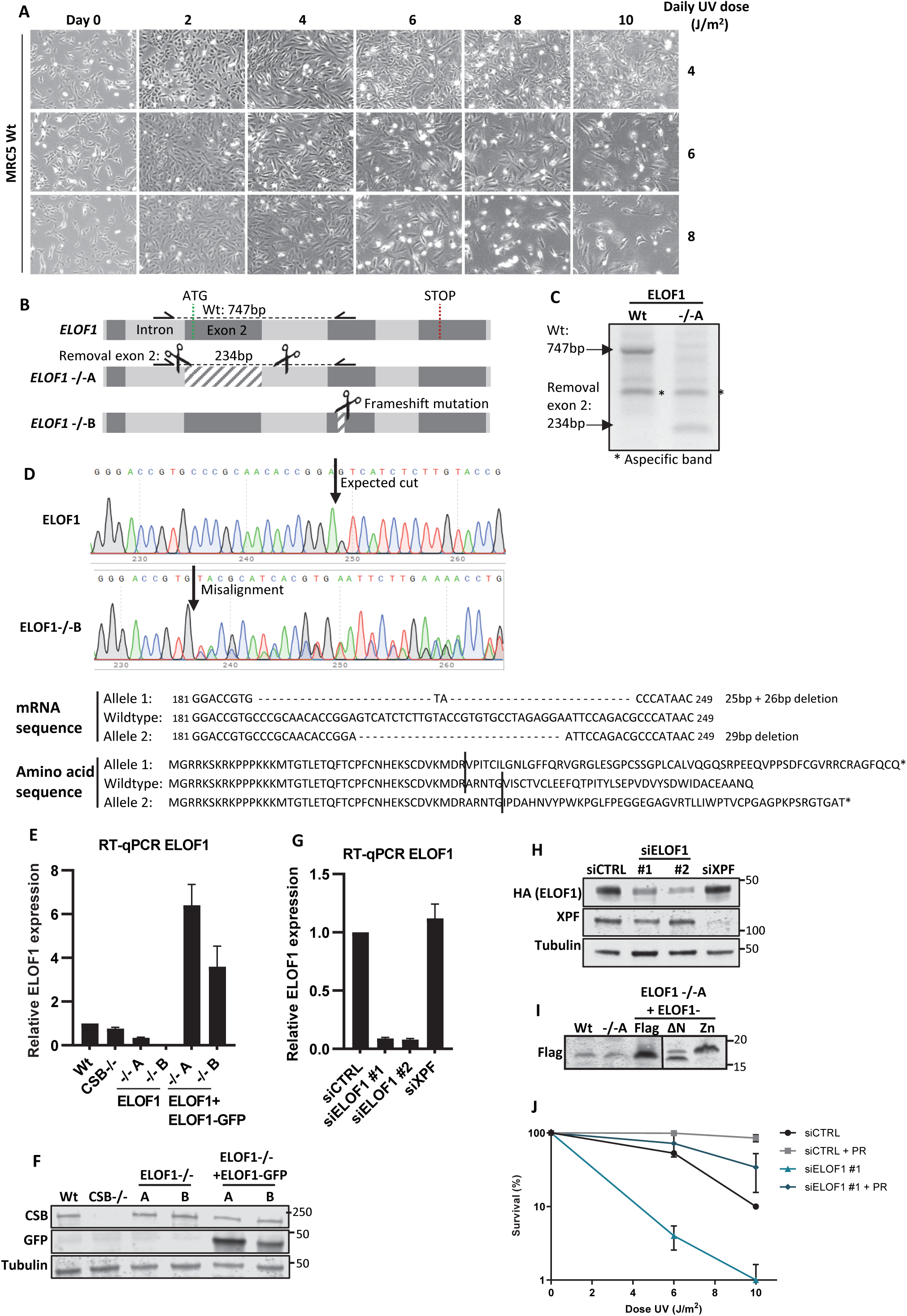
**(A)** Brightfield images of MRC-5 cells irradiated with indicated doses of UV-C for 10 consecutive days. Images were taken every other day. **(B)** Schematic of the genomic *ELOF1* locus. Scissors indicate target regions of the sgRNAs used to generated *ELOF1* KO (-/-) cells, half arrows indicate primers used for genotyping as shown in (C). **(C+D)** Genotyping of ELOF1 KO (-/-) cells, both originating from a single cell clone. **(C)** Genotyping PCR of loss of exon 2 in *ELOF1* -/-A cells. **(D)** Top panel: Sequencing results showing frameshift mutations in the targeted genomic locus of *ELOF1* -/-B. Bottom panel: Amino acid sequence of ELOF1 in ELOF1 -/-B cells. **(E)** Relative ELOF1 levels in indicated HCT116 Wt and ELOF1 KO (-/-) cells, with ELOF1 re-expression where indicated, as determined by RT-qPCR. Relative ELOF1 mRNA expression was normalized to GAPDH signal and levels in Wt cells were set to 1. Error bars indicate SEM. **(F)** Immunoblot of indicated HCT116 cell lines showing CSB or ELOF1-GFP expression. Tubulin was used as loading control. **(G)** Relative ELOF1 levels in HCT116 cells transfected with indicated siRNAs as determined by RT-qPCR. Relative ELOF1 expression was normalized to GAPDH signal and siCTRL levels were set to 1. Error bars indicate SEM. **(H)** Immunoblot showing endogenous ELOF1 and XPF levels in *ELOF1-mScarletI-HA* KI cells (suppl. fig. 2A) transfected with indicated siRNAs. Tubulin was used as loading control. **(I)** Immunoblot showing expression of Flag-tagged Wt or indicated ELOF1 mutants in HCT116 ELOF1 -/-A cells. **(J)** Relative colony survival of CPD photolyase cells transfected with indicated siRNAs. PR indicates CPD removal by photoreactivation. Plotted curves represent averages of 2 independent experiments ± SEM.

**Supplemental figure 2.**
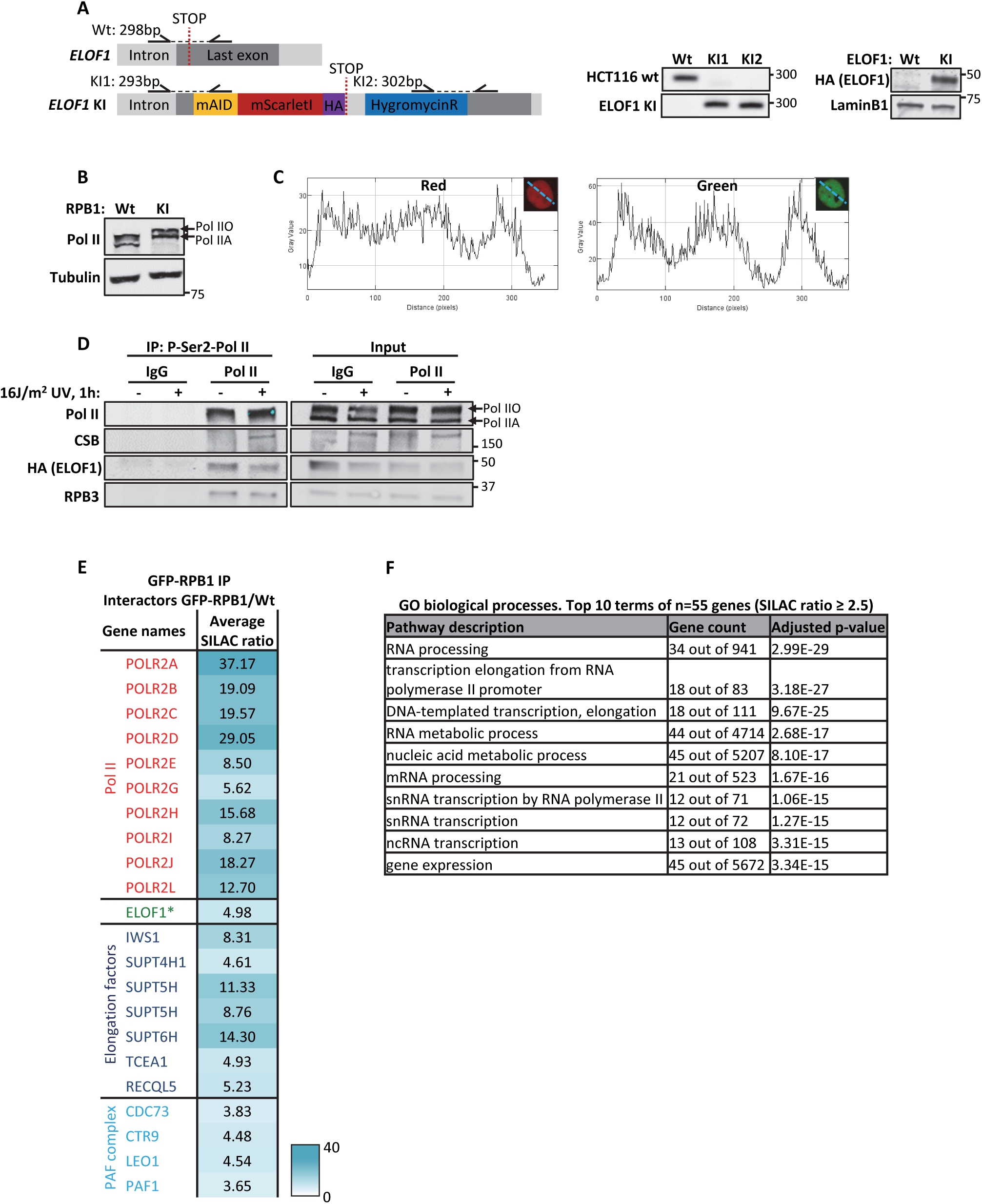
**(A)** Left panel: Schematic of the genomic locus of *ELOF1* for generating *ELOF1-mScarletI-HA* KI cell line. Half arrows indicate primer locations. Middle and right panel: Genotyping PCR and immunoblot for *ELOF1*-KI cell line. LaminB1 was used as loading control. **(B)** Immunoblot of HCT116 *GFP-RPB1* KI. Tubulin was used as loading control. **(C)** Histograms showing intensities of GFP and mScarletI measured over the indicated dotted line in HCT116 double KI cells. **(D)** Native immunoprecipitation of P-Ser2-modified Pol II in HCT116 cells followed by immunoblotting for indicated proteins. Cells were harvested 1 hour after mock treated or irradiation with 16 J/m^2^ UV-C. IgG was used as binding control. **(E)** Interaction heat map based on the SILAC ratios of MRC-5 GFP-RPB1-interacting proteins as determined by quantitative interaction proteomics. Average SILAC ratios of duplicate experiments are plotted and represent RPB1-interactors relative to empty beads. SILAC ratio >1 indicates increase in interaction. * indicates proteins quantified in one experiment. **(F)** Top 10 enriched GO terms (biological processes) identified using g:Profiler of 55 proteins identified as ELOF1 interactor with an average SILAC ratio of 2.5 or higher.

**Supplemental figure 3.**
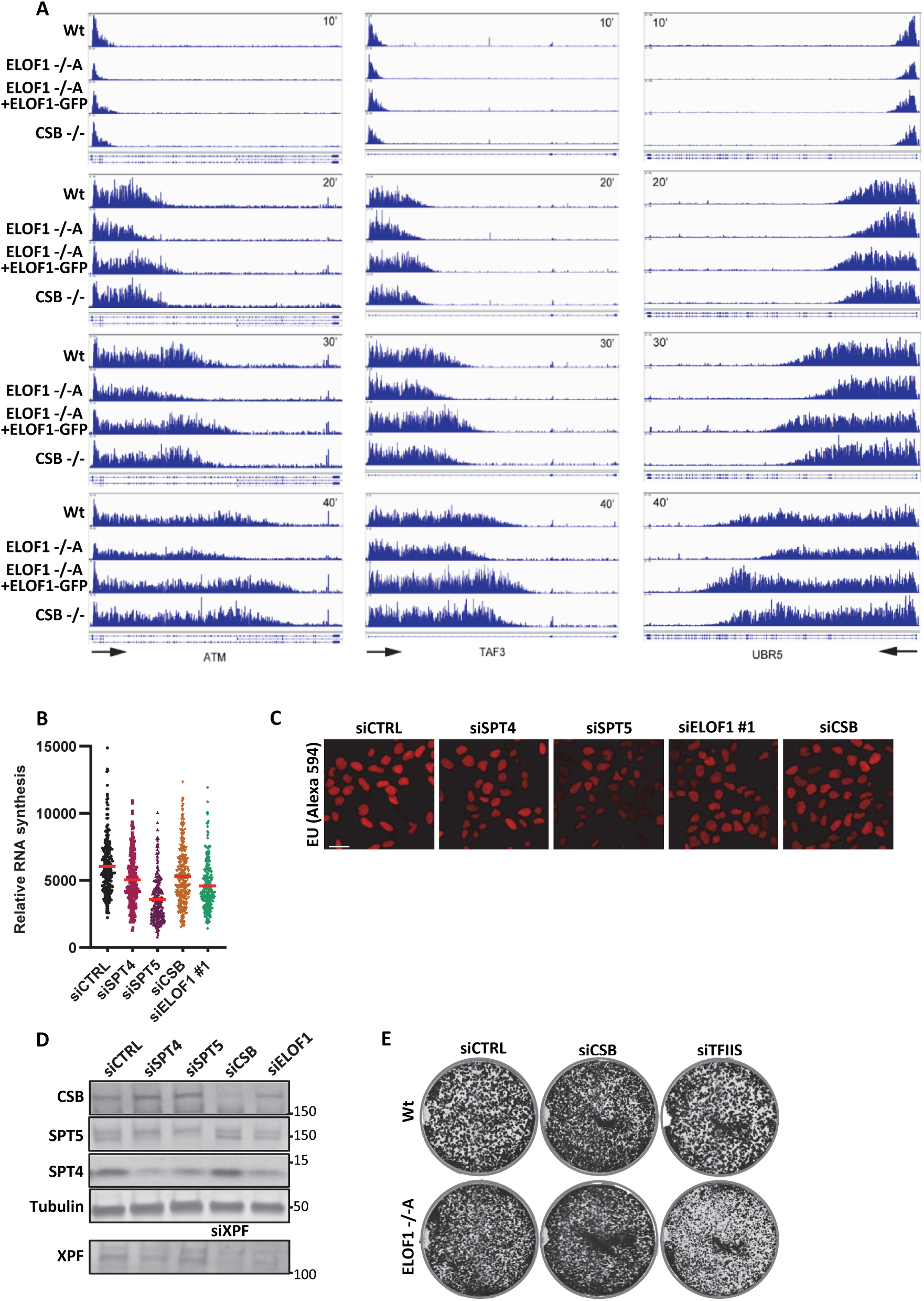
**(A)** Browser tracks from DRB/TT_chem_-seq experiment at *ATM, TAF3* and *UBR5*. Results are shown 10, 20, 30 or 40 minutes after DRB release. **(B)** Transcription levels as determined by relative EU incorporation in HCT116 cells transfected with indicated siRNAs. Red lines indicate average integrated density ± SEM. n≥200 cells from two independent experiments. **(C)** Representative images of EU incorporation in HCT116 cells transfected with indicated siRNAs. Scale bar: 20 µm. **(D)** Immunoblot for indicated proteins in HCT116 cells transfected with indicated siRNAs. Tubulin was used as loading control. **(E)** Images of HCT116 Wt and ELOF1 -/-A cells transfected with indicated siRNAs, stained with coomassie blue 10 days after transfection.

**Supplemental figure 4.**
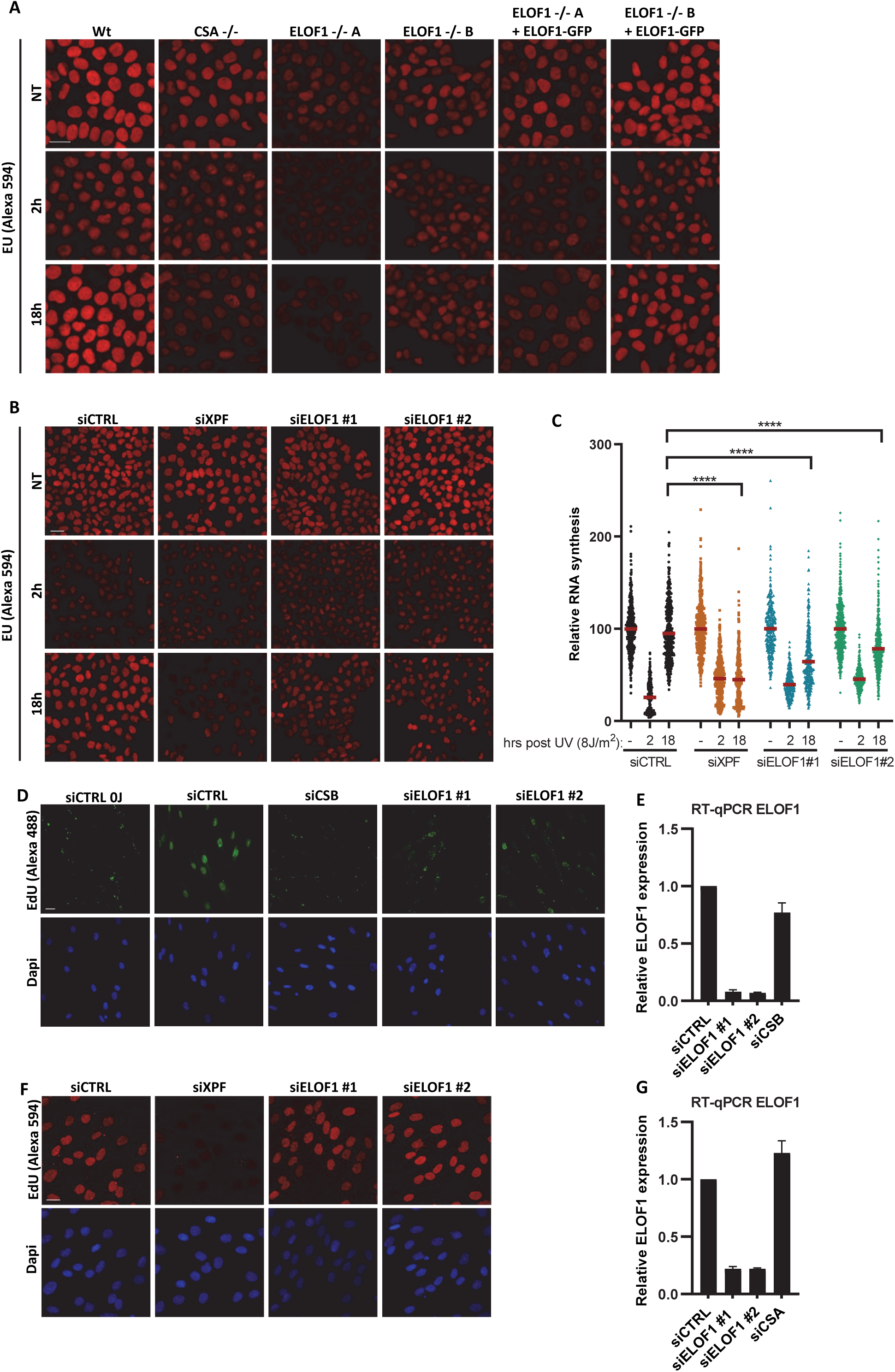
**(A+B)** Representative immunofluorescence images of EU incorporation in **(A)** indicated HCT116 Wt and KO (-/-) cells, with ELOF1 re-expression where indicated, or **(B)** HCT116 cells transfected with indicated siRNAs, 2 or 18 hours after 8 J/m^2^ UV-C or mock treatment (NT). Scale bar: 20 µm. **(C)** Transcription restart after UV damage as determined by relative EU incorporation in HCT116 cells transfected with indicated siRNAs, 2 or 18 hours after 8 J/m^2^ UV-C or mock treatment (NT). Relative integrated density of UV-irradiated samples are normalized to mock-treated and set to 100%. Red lines indicate average integrated density ± SEM. n≥300 cells from three independent experiments. **(D)** Representative immunofluorescence images of amplified EdU signal in XP186LV fibroblasts (XP-C) transfected with indicated siRNAs, 7 hours after exposure to 8 J/m^2^ UV-C. Scalebar: 20 µm. **(E)** Relative ELOF1 mRNA levels in XP186LV fibroblasts (XP-C) following transfection with indicated siRNAs as determined by RT-qPCR. ELOF1 expression was normalized to GAPDH expression and siCTRL levels were set to 1. Error bars indicate SEM. **(F)** Representative fluorescence images of EdU incorporation 3 hours after irradiation with 16 J/m^2^ UV-C in C5RO (hTert) cells transfected with indicated siRNAs. Scale bar: 20 µm. **(G)** Relative ELOF1 mRNA levels in C5RO (hTert) cells following transfection with indicated siRNAs as determined by RT-qPCR. ELOF1 expression was normalized to GAPDH expression and siCTRL levels were set to 1. Error bars indicate SEM. ****P ≤0.0001.

**Supplemental figure 5.**
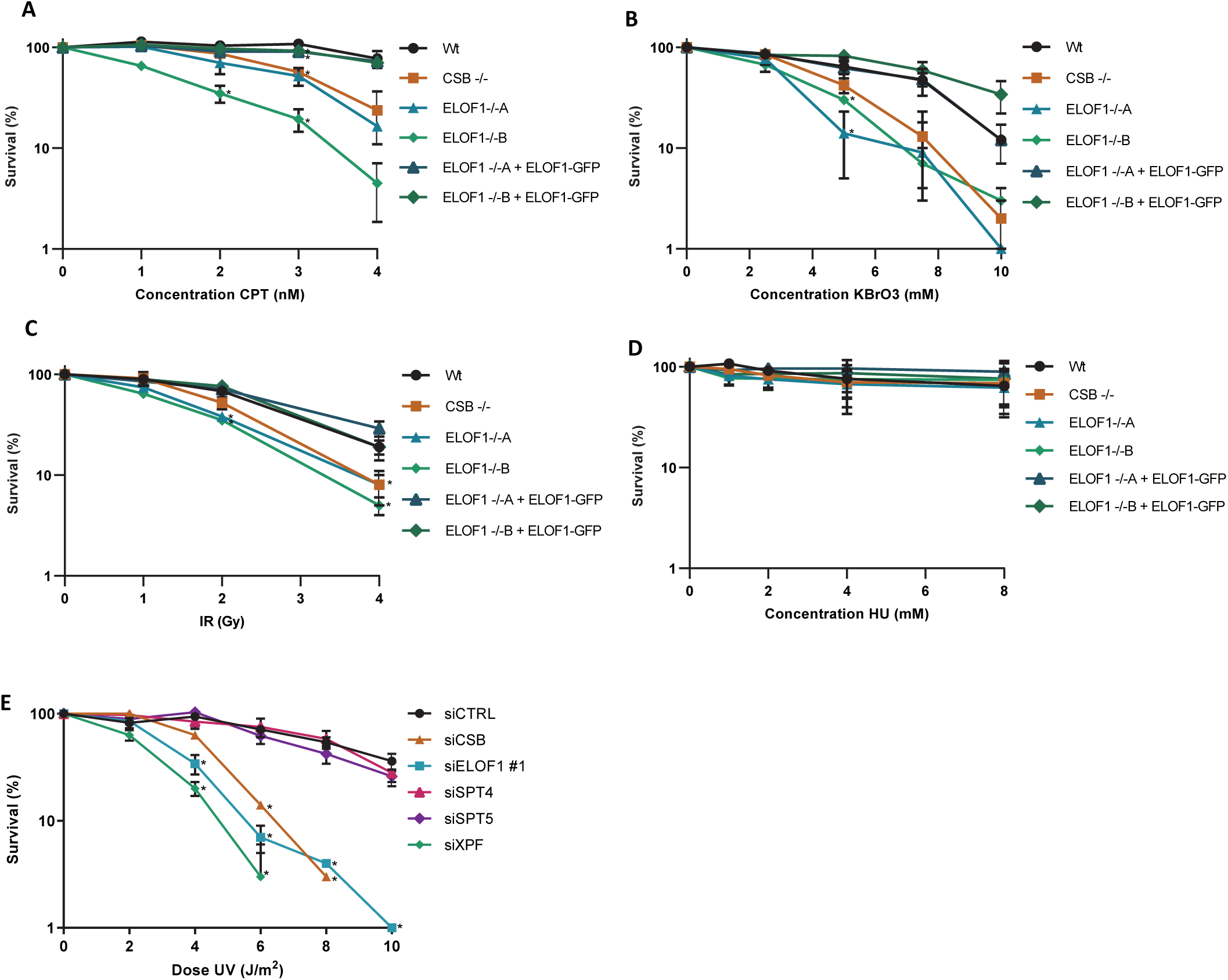
**(A-D)** Relative colony survival of indicated HCT116 Wt and KO (-/-) cells, with ELOF1 re-expression where indicated, continuously exposed to indicated concentrations of **(A)** camptothecin (CPT) or **(B)** potassium bromate (KBrO_3_), or irradiated with indicated doses of **(C)** ionizing radiation (IR), or exposed **(D)** to indicated concentrations of hydroxyurea (HU). Plotted curves represent averages of at least two independent experiments ± SEM. **(E)** Relative colony survival of HCT116 cells transfected with indicated siRNAs following exposure to indicated doses of UV-C. Plotted curves represent averages of three independent experiments ± SEM. *P≤0.05

**Supplemental figure 6.**
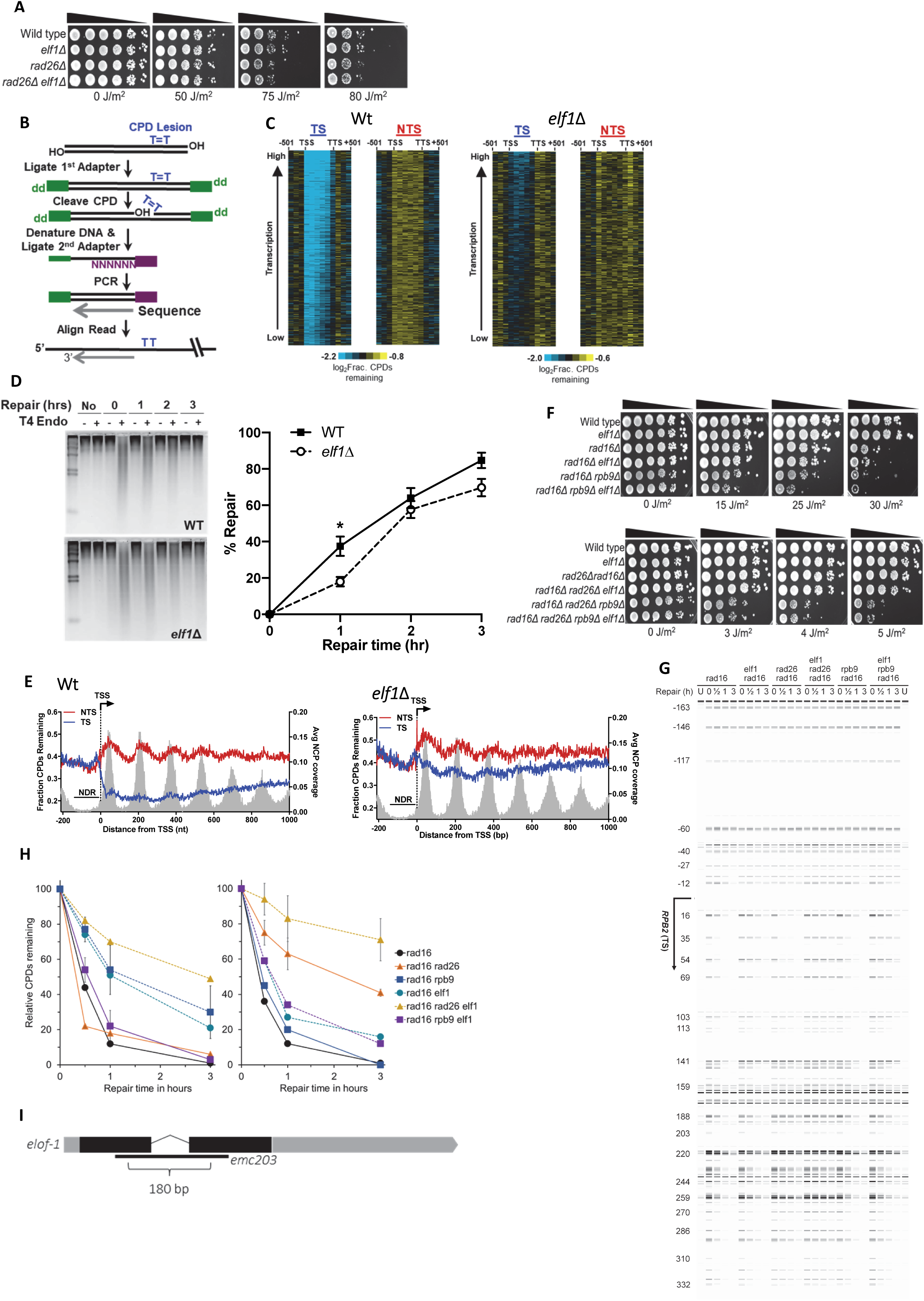
**(A)** Indicated mutant yeast strains were serially 10-fold diluted, spotted, and exposed to indicated UV-C doses. **(B)** Schematic showing the CPD-seq method. Isolated DNA is sonicated and adaptors are ligated. Subsequently, CPDs are cleaved by T4 endonuclease V and APE1 nuclease to generate 3’ ends. Following denaturing of the DNA, the ends are ligated to a second adaptor that allows sequencing of CPDs. **(C)** Gene plot analysis of CPD-seq data following 2-hour repair for ∼4500 yeast genes, ordered by transcription frequency. Plots depict fraction of unrepaired CPDs following 2-hour repair relative to no repair for both the transcribed strand (TS) and non-transcribed strand (NTS) for gene coding regions, regions upstream of the transcription start site (TSS), and downstream of the transcription termination site (TTS). Each row represents approximately 10 genes to display the plot in a compact manner. **(D)** Analysis of bulk repair of UV-induced CPD lesions in Wt and *elf1*Δ mutant yeast. The repair of CPD lesions at various time points was measured by T4 endonuclease V digestion and alkaline gel electrophoresis of genomic DNA isolated from UV-irradiated yeast (100 J/m^2^ UV-C light). A representative gel is shown on the left. The right panel depicts the quantification of CPD repair at each time point from at least three independent experiments ±SEM. \**P*≤0.05. **(E)** Single nucleotide resolution analysis of CPD-seq data downstream of the TTS of ∼4500 yeast genes. Plots depict fraction of unrepaired CPDs following 2-hour repair relative to no repair for both TS and NTS. Nucleosome positioning data is shown for reference. **(F)** Controls for UV spotting assays shown in Fig. 3I. **(G)** Image showing repair of CPDs in the TS of the *RPB2* gene for indicated yeast strains. The image was generated by converting counts of sequencing reads aligned to the sites of the RPB2 fragment into bands. ‘*U*’ indicates samples from unirradiated cells. Nucleotide positions relative to the TSS (+1) of the *RPB2* gene are indicated on the left. **(H)** Left panel: Relative percentage of CPDs remaining in the short region (within 54 bp) immediately downstream of the transcription start site of the *RPB2* gene. Right panel: Relative percentage of CPDs remaining in the more downstream region (from 69 to 353 bp) of the *RPB2* gene. Error bars (S.D.) are shown only for most relevant strains for clarity. **(I)** Schematic representation of the *C. elegans elof-1* genomic organization, depicting the 180 bp *emc203* deletion allele generated with CRISPR/Cas9. Shaded boxes represent exons with coding sequences shown in black.

**Supplemental figure 7.**
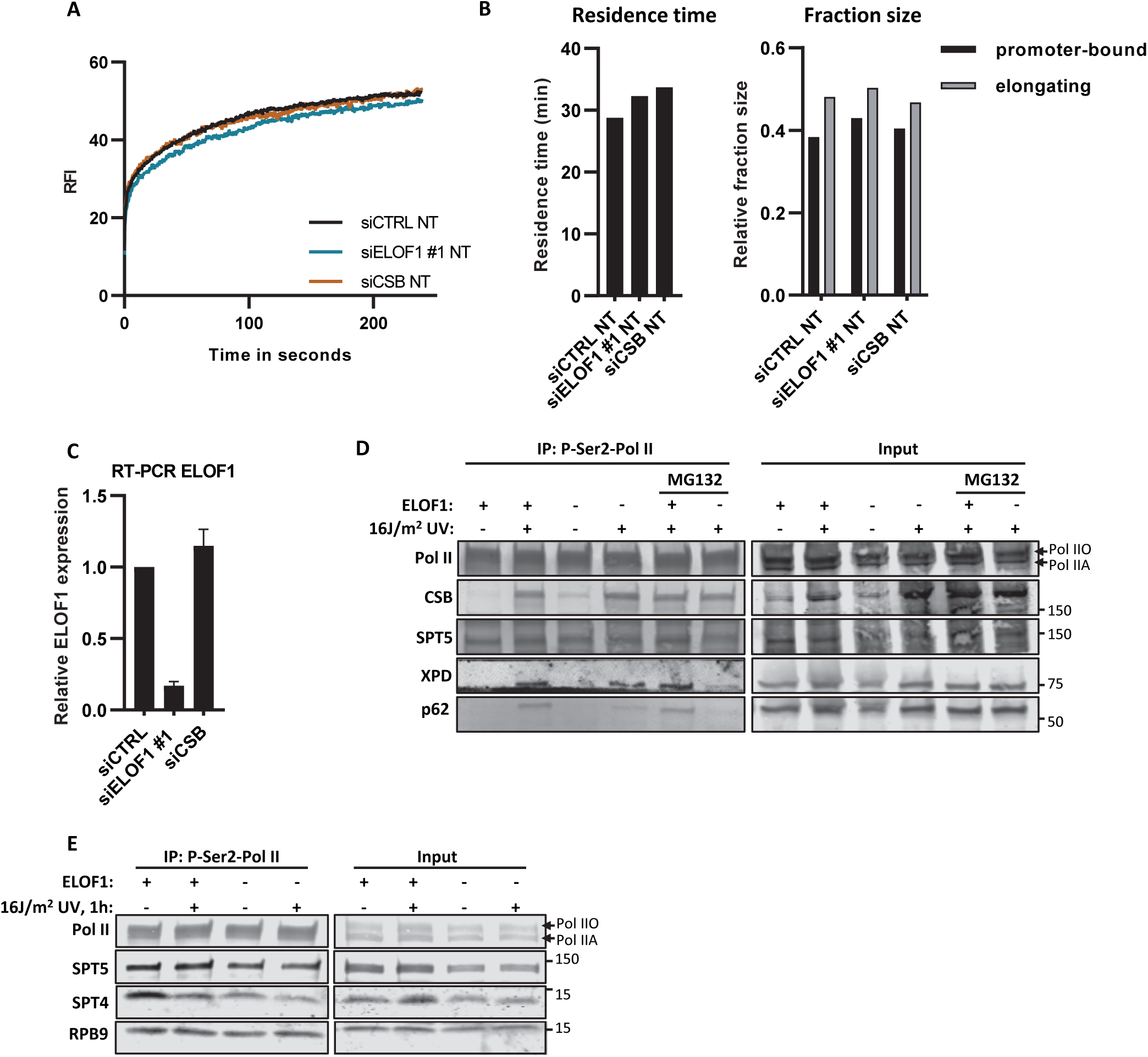
**(A)** FRAP analysis of GFP-RPB1 mobility after depletion of indicated factors. Mock-treated curves corresponding to figure 4A. n≥14 cells. **(B)** Left panel: Residence time of elongating Pol II or right panel: relative fraction size of promoter-bound or elongating Pol II as determined by Monte-Carlo-based modeling of RPB1 mobility as shown in (A). **(C)** Relative ELOF1 mRNA levels in *GFP-RPB1* KI cells transfected with indicated siRNAs as determined by RT-qPCR. ELOF1 expression was normalized to GAPDH signal and levels of control cells were set to 1. Error bars indicate SEM. **(D)** Native immunoprecipitation of Pol II in Wt and ELOF -/-A cells followed by immunoblotting for indicated proteins. Cells were harvested 1 hour after mock treatment or irradiation with 16 J/m^2^ UV-C. MG132: treatment with 50 µM proteasome inhibitor MG132, 1 hour before UV irradiation. **(E)** Native immunoprecipitation of Pol II in Wt and ELOF -/-A cells followed by immunoblotting for indicated proteins. Cells were harvested 1 hour after mock treatment or irradiation with 16 J/m^2^ UV-C.

**Supplemental figure 8.**
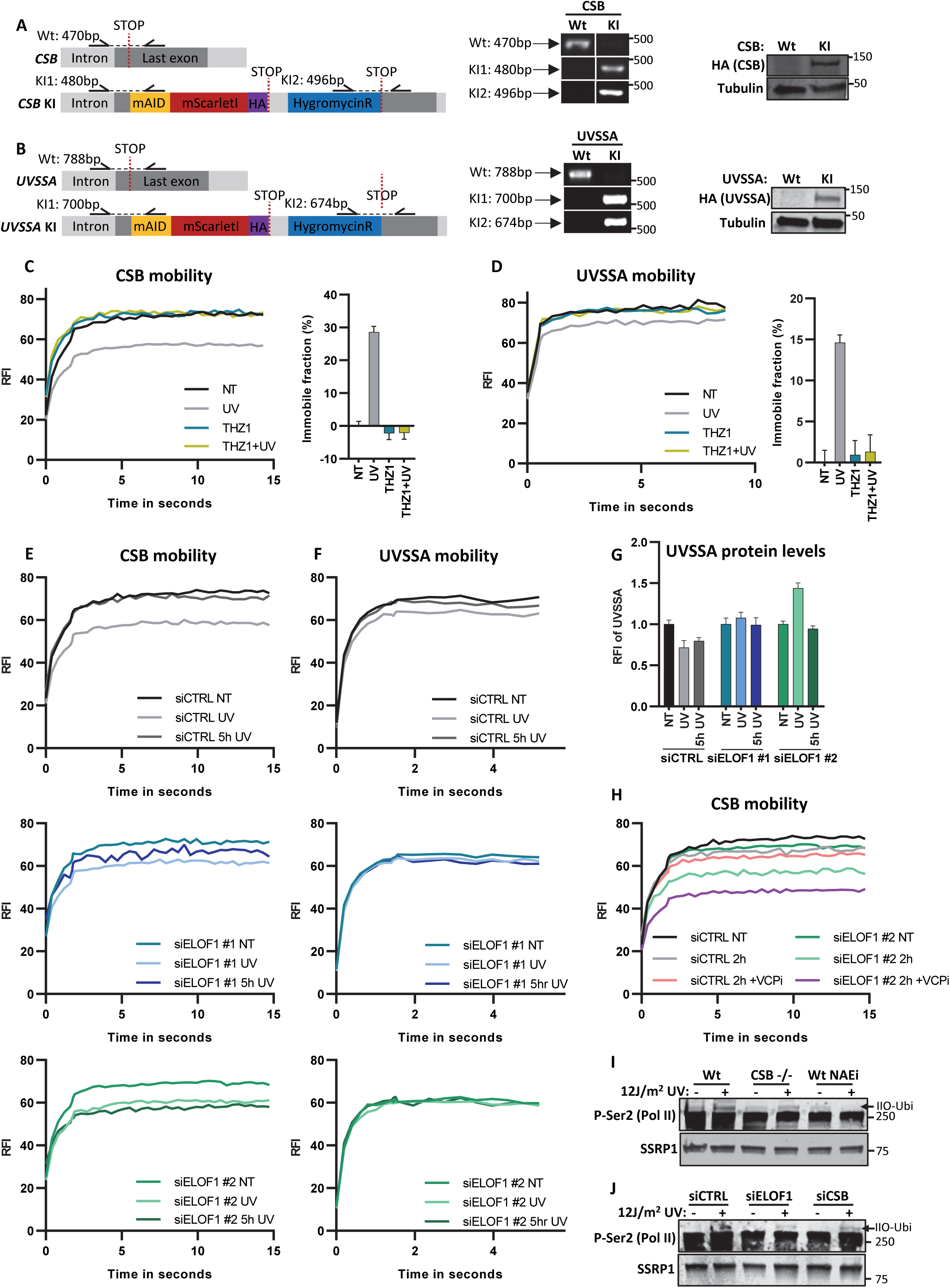
**(A)** Left panel: Schematic of the genomic locus of *CSB* and used strategy for generating the homozygous *CSB-mScarletI-HA* KI cell line. Half arrows indicate primer locations. Middle and right panel: Genotyping PCR and immunoblot for *CSB*-KI cell line. **(B)** Left panel: Schematic of the genomic locus of *UVSSA* and used strategy for generating the homozygous *UVSSA-mScarletI-HA* KI cell line. Half arrows indicate primer locations. Middle and right panel: Genotyping PCR and immunoblot for *UVSSA*-KI cell line. **(C)** Left panel: CSB mobility was determined by FRAP analysis of CSB-mScarletI after the indicated treatments. THZ1: 1 hour treatment (2 µM) before UV-C irradiation (4 J/m^2^) or mock treatment. Right panel: Relative immobile fraction of CSB as determined by FRAP analysis. Plotted values represent mean ± SEM and are normalized to mock treated. n≥15 cells. **(D)** Same as C but for UVSSA-mScarletI. n≥10 cells. **(E+F)** FRAP analyses of CSB-mScarletI **(E)** or UVSSA-mScarletI **(F)** mobility after transfection with indicated siRNAs in individual graphs. Cells were mock treated (NT) or analyzed directly (UV) or 5 hours (5hr UV) after irradiation with 4 J/m^2^ UV-C. **(G)** Relative fluorescence intensity of UVSSA in *UVSSA*-KI cells transfected with indicated siRNAs as determined by live-cell imaging. Plotted values represent mean ± SEM. n≥16 cells. **(H)** FRAP analysis of CSB in *CSB*-KI cells transfected with indicated siRNAs 2 hours after UV. VCPi: VCP inhibitor (5 µM) was directly added after UV-C (4 J/m^2^). **(I)** Immunoblot of chromatin fraction of indicated cell lines 1 hour after 12 J/m^2^ UV-C or mock treatment. NAEi = 1 hour treatment with NEDDylation inhibitor (10 µM). SSRP1 is shown as loading control. **(J)** Immunoblot of chromatin fraction of HCT116 cells transfected with indicated siRNAs 1 hour after 12 J/m^2^ UV-C or mock treatment. SSRP1 is shown as loading control.

**Supplemental figure 9.**
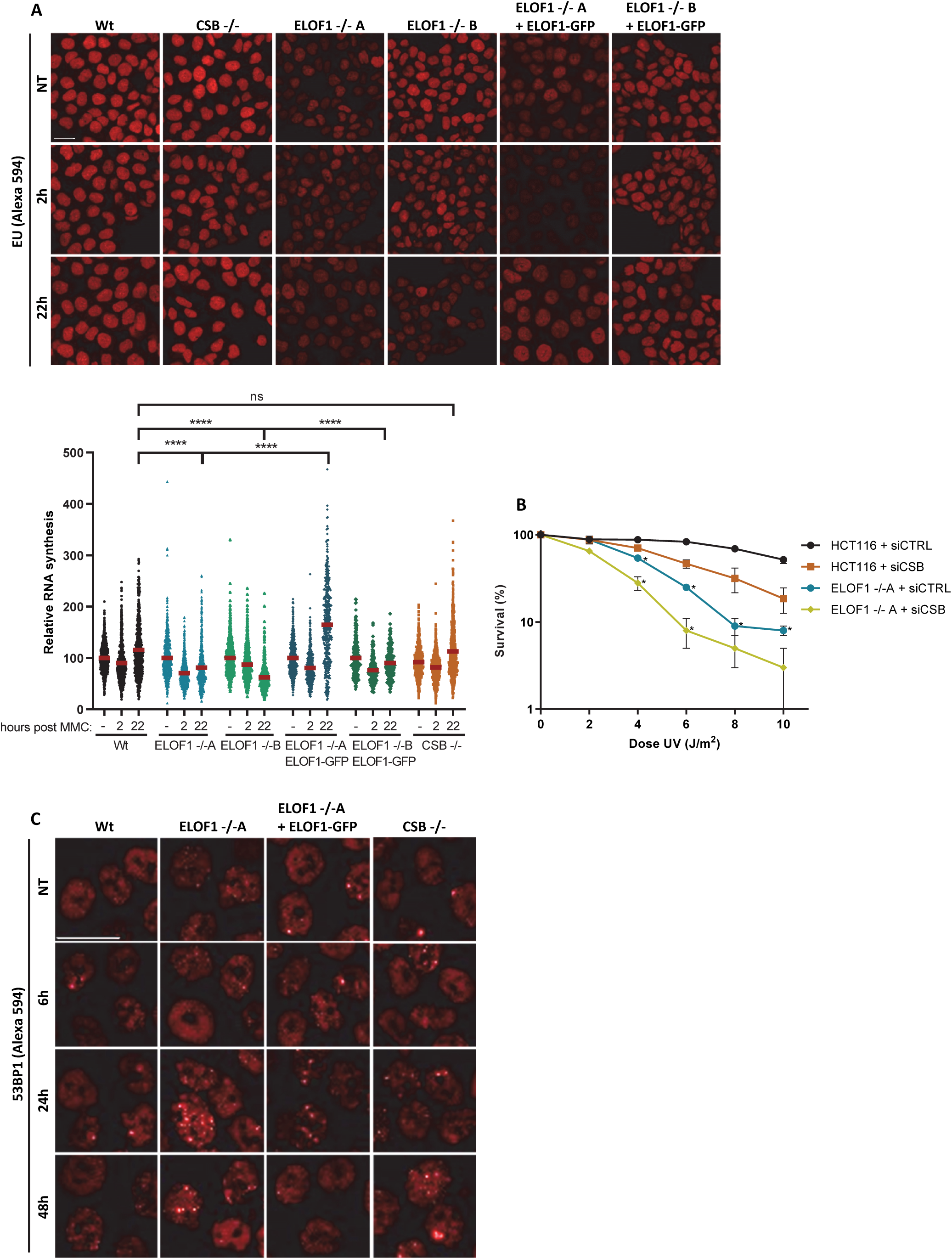
**(A)** Top panel: Representative immunofluorescence images of EU incorporation in indicated HCT116 Wt and KO (-/-) cells, with ELOF1 re-expression where indicated, 2 or 22 hours after a 2-hour exposure to 10 µg/ml mitomycin C or mock treatment. Scale bar: 20 µm. Bottom panel: Transcription restart after mitomycin C as determined by relative EU incorporation in the indicated HCT116 cells. Mitomycin C-treated samples are normalized to mock treated levels and set to 100%. Red lines indicate average integrated density ± SEM. n≥300 cells from four independent experiments. **(B)** Relative colony survival of indicated cell lines with siRNA transfection following exposure to indicated doses of UV-C. Plotted curves represent averages of three independent experiments ± SEM. **(C)** Representative immunofluorescence images of 53BP1 foci in indicated HCT116 Wt and KO (-/-) cells, with ELOF1 re-expression where indicated, 6, 24 or 48 hours after exposure to 8 J/m^2^ UV-C or mock treatment. Scale bar: 20 µm. *p≤0.05, ****p≤0.0001.

**Supplemental figure 10.**
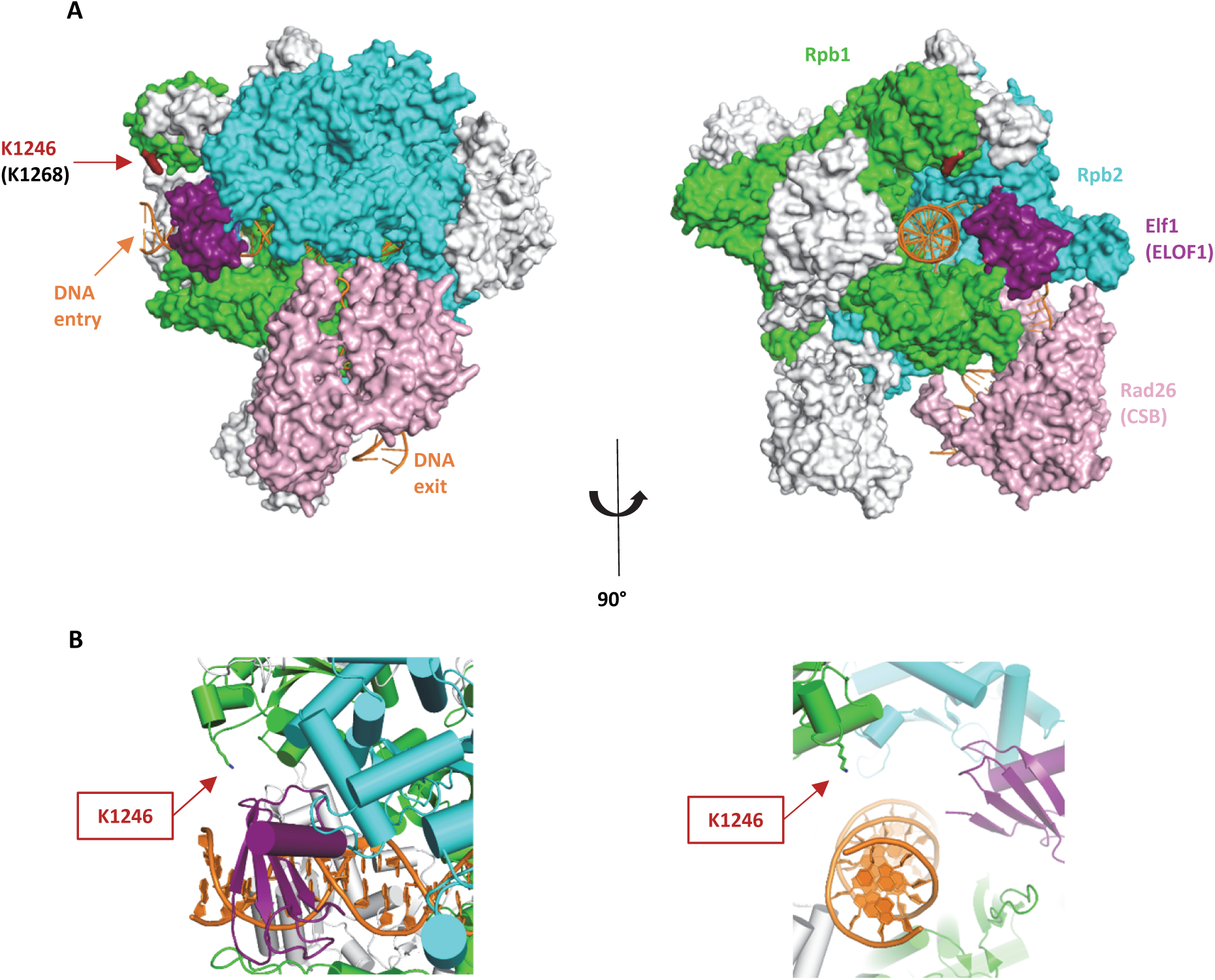
**(A)** *S.cerevisiae* Pol II (5vvr.pdb) with Rpb1 in green, Rpb2 in cyan, DNA in orange and Rad26 (CSB) in pink. The *P.pastoris* Pol II in complex with elongation factors (5xog.pdb) was superimposed onto this structure (Rpb1 subunits aligned onto each other), and all subunits except Elf1 (ELOF1; purple) were omitted for clarity. Conserved lysine K1246 (K1268 in mammalian Pol II) is indicated in dark red. **(B)** Close up of Elf1 (ELOF1) binding region.

**Supplemental table 1.** Table showing negatively regulated genes from the CRISPR/cas9 screen resulting from MaGecK analysis of the change in abundance of sgRNAs in UV-treated over mock-treated samples. Experiment was performed in duplicate.

**Supplemental table 2.** Table with SILAC ratios and peptide numbers as determined using quantitative interaction proteomics. Each tab represents a different experiment as indicated.

## References

1. Lans, H., Hoeijmakers, J.H.J., Vermeulen, W. & Marteijn, J.A. The DNA damage response to transcription stress. Nat Rev Mol Cell Biol 20, 766–784 (2019).

2. Gregersen, L.H. & Svejstrup, J.Q. The Cellular Response to Transcription-Blocking DNA Damage. Trends Biochem Sci 43, 327–341 (2018).

3. Doetsch, P.W. Translesion synthesis by RNA polymerases: occurrence and biological implications for transcriptional mutagenesis. Mutat Res 510, 131–40 (2002).

4. Marietta, C. & Brooks, P.J. Transcriptional bypass of bulky DNA lesions causes new mutant RNA transcripts in human cells. EMBO Rep 8, 388–93 (2007).

5. Hoeijmakers, J.H. DNA damage, aging, and cancer. N Engl J Med 361, 1475–85 (2009).

6. Helmrich, A., Ballarino, M., Nudler, E. & Tora, L. Transcription-replication encounters, consequences and genomic instability. Nat Struct Mol Biol 20, 412–8 (2013).

7. Ljungman, M. & Zhang, F. Blockage of RNA polymerase as a possible trigger for u.v. light-induced apoptosis. Oncogene 13, 823–31 (1996).

8. Gomez-Gonzalez, B. & Aguilera, A. Transcription-mediated replication hindrance: a major driver of genome instability. Genes Dev 33, 1008–1026 (2019).

9. Crossley, M.P., Bocek, M. & Cimprich, K.A. R-Loops as Cellular Regulators and Genomic Threats. Mol Cell 73, 398–411 (2019).

10. Gaillard, H. & Aguilera, A. Transcription as a Threat to Genome Integrity. Annu Rev Biochem 85, 291–317 (2016).

11. Hanawalt, P.C. & Spivak, G. Transcription-coupled DNA repair: two decades of progress and surprises. Nat Rev Mol Cell Biol 9, 958–70 (2008).

12. Laugel, V. Cockayne syndrome: the expanding clinical and mutational spectrum. Mech Ageing Dev 134, 161–70 (2013).

13. Xu, J. et al. Structural basis for the initiation of eukaryotic transcription-coupled DNA repair. Nature 551, 653–657 (2017).

14. Saijo, M. et al. Functional TFIIH is required for UV-induced translocation of CSA to the nuclear matrix. Mol Cell Biol 27, 2538–47 (2007).

15. Kamiuchi, S. et al. Translocation of Cockayne syndrome group A protein to the nuclear matrix: possible relevance to transcription-coupled DNA repair. Proc Natl Acad Sci U S A 99, 201–6 (2002).

16. Sin, Y., Tanaka, K. & Saijo, M. The C-terminal Region and SUMOylation of Cockayne Syndrome Group B Protein Play Critical Roles in Transcription-coupled Nucleotide Excision Repair. J Biol Chem 291, 1387–97 (2016).

17. Groisman, R. et al. CSA-dependent degradation of CSB by the ubiquitin-proteasome pathway establishes a link between complementation factors of the Cockayne syndrome. Genes Dev 20, 1429–34 (2006).

18. Nakazawa, Y. et al. Ubiquitination of DNA Damage-Stalled RNAPII Promotes Transcription-Coupled Repair. Cell 180, 1228–1244 e24 (2020).

19. Tufegdzic Vidakovic, A. et al. Regulation of the RNAPII Pool Is Integral to the DNA Damage Response. Cell 180, 1245–1261 e21 (2020).

20. Okuda, M., Nakazawa, Y., Guo, C., Ogi, T. & Nishimura, Y. Common TFIIH recruitment mechanism in global genome and transcription-coupled repair subpathways. Nucleic Acids Res 45, 13043–13055 (2017).

21. Schärer, O.D. Nucleotide excision repair in eukaryotes. Cold Spring Harb Perspect Biol 5, a012609 (2013).

22. Marteijn, J.A., Lans, H., Vermeulen, W. & Hoeijmakers, J.H. Understanding nucleotide excision repair and its roles in cancer and ageing. Nat Rev Mol Cell Biol 15, 465–81 (2014).

23. Sanjana, N.E., Shalem, O. & Zhang, F. Improved vectors and genome-wide libraries for CRISPR screening. Nat Methods 11, 783–784 (2014).

24. Evers, B. et al. CRISPR knockout screening outperforms shRNA and CRISPRi in identifying essential genes. Nat Biotechnol 34, 631–3 (2016).

25. Li, W. et al. MAGeCK enables robust identification of essential genes from genome-scale CRISPR/Cas9 knockout screens. Genome Biol 15, 554 (2014).

26. Yang, W. & Gao, Y. Translesion and Repair DNA Polymerases: Diverse Structure and Mechanism. Annu Rev Biochem 87, 239–261 (2018).

27. Daniels, J.P., Kelly, S., Wickstead, B. & Gull, K. Identification of a crenarchaeal orthologue of Elf1: implications for chromatin and transcription in Archaea. Biol Direct 4, 24 (2009).

28. Prather, D., Krogan, N.J., Emili, A., Greenblatt, J.F. & Winston, F. Identification and characterization of Elf1, a conserved transcription elongation factor in Saccharomyces cerevisiae. Mol Cell Biol 25, 10122–35 (2005).

29. Joo, Y.J., Ficarro, S.B., Chun, Y., Marto, J.A. & Buratowski, S. In vitro analysis of RNA polymerase II elongation complex dynamics. Genes Dev 33, 578–589 (2019).

30. Ehara, H. et al. Structure of the complete elongation complex of RNA polymerase II with basal factors. Science 357, 921–924 (2017).

31. Mayer, A. et al. Uniform transitions of the general RNA polymerase II transcription complex. Nat Struct Mol Biol 17, 1272–8 (2010).

32. Ehara, H. et al. Structural insight into nucleosome transcription by RNA polymerase II with elongation factors. Science 363, 744–747 (2019).

33. Steurer, B. et al. Live-cell analysis of endogenous GFP-RPB1 uncovers rapid turnover of initiating and promoter-paused RNA Polymerase II. Proc Natl Acad Sci U S A 115, E4368–E4376 (2018).

34. Nilson, K.A. et al. THZ1 Reveals Roles for Cdk7 in Co-transcriptional Capping and Pausing. Mol Cell 59, 576–87 (2015).

35. Chao, S.H. et al. Flavopiridol inhibits P-TEFb and blocks HIV-1 replication. J Biol Chem 275, 28345–8 (2000).

36. Sobell, H.M. Actinomycin and DNA transcription. Proc Natl Acad Sci U S A 82, 5328–31 (1985).

37. Van Oss, S.B., Cucinotta, C.E. & Arndt, K.M. Emerging Insights into the Roles of the Paf1 Complex in Gene Regulation. Trends Biochem Sci 42, 788–798 (2017).

38. Chen, F.X., Smith, E.R. & Shilatifard, A. Born to run: control of transcription elongation by RNA polymerase II. Nat Rev Mol Cell Biol 19, 464–478 (2018).

39. van den Boom, V. et al. DNA damage stabilizes interaction of CSB with the transcription elongation machinery. J Cell Biol 166, 27–36 (2004).

40. Gregersen, L.H., Mitter, R. & Svejstrup, J.Q. Using TTchem-seq for profiling nascent transcription and measuring transcript elongation. Nat Protoc 15, 604–627 (2020).

41. Nudler, E. RNA polymerase backtracking in gene regulation and genome instability. Cell 149, 1438–45 (2012).

42. Jia, N. et al. A rapid, comprehensive system for assaying DNA repair activity and cytotoxic effects of DNA-damaging reagents. Nat Protoc 10, 12–24 (2015).

43. Wienholz, F., Vermeulen, W. & Marteijn, J.A. Amplification of unscheduled DNA synthesis signal enables fluorescence-based single cell quantification of transcription-coupled nucleotide excision repair. Nucleic Acids Res 45, e68 (2017).

44. Jaspers, N.G. et al. Anti-tumour compounds illudin S and Irofulven induce DNA lesions ignored by global repair and exclusively processed by transcription- and replication-coupled repair pathways. DNA Repair (Amst) 1, 1027–38 (2002).

45. Slyskova, J. et al. Base and nucleotide excision repair facilitate resolution of platinum drugs-induced transcription blockage. Nucleic Acids Res 46, 9537–9549 (2018).

46. Veloso, A. et al. Genome-wide transcriptional effects of the anti-cancer agent camptothecin. PLoS One 8, e78190 (2013).

47. Brooks, P.J. et al. The oxidative DNA lesion 8,5’-(S)-cyclo-2’-deoxyadenosine is repaired by the nucleotide excision repair pathway and blocks gene expression in mammalian cells. J Biol Chem 275, 22355–62 (2000).

48. van Gool, A.J. et al. RAD26, the functional S. cerevisiae homolog of the Cockayne syndrome B gene ERCC6. EMBO J 13, 5361–9 (1994).

49. Mao, P., Smerdon, M.J., Roberts, S.A. & Wyrick, J.J. Chromosomal landscape of UV damage formation and repair at single-nucleotide resolution. Proc Natl Acad Sci U S A 113, 9057–62 (2016).

50. Tijsterman, M., Verhage, R.A., van de Putte, P., Tasseron-de Jong, J.G. & Brouwer, J. Transitions in the coupling of transcription and nucleotide excision repair within RNA polymerase II-transcribed genes of Saccharomyces cerevisiae. Proc Natl Acad Sci U S A 94, 8027–32 (1997).

51. Li, S. & Smerdon, M.J. Rpb4 and Rpb9 mediate subpathways of transcription-coupled DNA repair in Saccharomyces cerevisiae. EMBO J 21, 5921–9 (2002).

52. Lans, H. et al. Involvement of global genome repair, transcription coupled repair, and chromatin remodeling in UV DNA damage response changes during development. PLoS Genet 6, e1000941 (2010).

53. Geverts, B., van Royen, M.E. & Houtsmuller, A.B. Analysis of biomolecular dynamics by FRAP and computer simulation. Methods Mol Biol 1251, 109–33 (2015).

54. Williamson, L. et al. UV Irradiation Induces a Non-coding RNA that Functionally Opposes the Protein Encoded by the Same Gene. Cell 168, 843–855 e13 (2017).

55. Brueckner, F., Hennecke, U., Carell, T. & Cramer, P. CPD damage recognition by transcribing RNA polymerase II. Science 315, 859–62 (2007).

56. van der Weegen, Y. et al. The sequential and cooperative action of CSB, CSA and UVSSA targets the TFIIH complex to DNA damage-stalled RNA polymerase II. bioRxiv, 707216 (2019).

57. Schwertman, P. et al. UV-sensitive syndrome protein UVSSA recruits USP7 to regulate transcription-coupled repair. Nat Genet 44, 598–602 (2012).

58. Zhang, X. et al. Mutations in UVSSA cause UV-sensitive syndrome and destabilize ERCC6 in transcription-coupled DNA repair. Nat Genet 44, 593–7 (2012).

59. He, J., Zhu, Q., Wani, G., Sharma, N. & Wani, A.A. Valosin-containing Protein (VCP)/p97 Segregase Mediates Proteolytic Processing of Cockayne Syndrome Group B (CSB) in Damaged Chromatin. J Biol Chem 291, 7396–408 (2016).

60. Lukas, C. et al. 53BP1 nuclear bodies form around DNA lesions generated by mitotic transmission of chromosomes under replication stress. Nat Cell Biol 13, 243–53 (2011).

61. Tellier, A.P., Archambault, D., Tremblay, K.D. & Mager, J. The elongation factor Elof1 is required for mammalian gastrulation. PLoS One 14, e0219410 (2019).

62. Fei, J. & Chen, J. KIAA 1530 protein is recruited by Cockayne syndrome complementation group protein A (CSA) to participate in transcription-coupled repair (TCR). J Biol Chem 287, 35118–26 (2012).

63. Wienholz, F. et al. FACT subunit Spt16 controls UVSSA recruitment to lesion-stalled RNA Pol II and stimulates TC-NER. Nucleic Acids Res (2019).

64. Poli, J. et al. Mec1, INO80, and the PAF1 complex cooperate to limit transcription replication conflicts through RNAPII removal during replication stress. Genes Dev 30, 337–54 (2016).

65. Campeau, E. et al. A Versatile Viral System for Expression and Depletion of Proteins in Mammalian Cells. PLOS ONE 4, e6529 (2009).

66. Yesbolatova, A., Natsume, T., Hayashi, K.-i. & Kanemaki, M.T. Generation of conditional auxin-inducible degron (AID) cells and tight control of degron-fused proteins using the degradation inhibitor auxinole. Methods 164–165, 73-80 (2019).

67. Natsume, T., Kiyomitsu, T., Saga, Y. & Kanemaki, M.T. Rapid Protein Depletion in Human Cells by Auxin-Inducible Degron Tagging with Short Homology Donors. Cell Rep 15, 210–218 (2016).

68. Steurer, B. et al. Live-cell analysis of endogenous GFP-RPB1 uncovers rapid turnover of initiating and promoter-paused RNA Polymerase II. Proceedings of the National Academy of Sciences 115, E4368–E4376 (2018).

69. Steurer, B. et al. Fluorescently-labelled CPD and 6-4PP photolyases: new tools for live-cell DNA damage quantification and laser-assisted repair. Nucleic Acids Research 47, 3536–3549 (2019).

70. Brinkman, E.K., Chen, T., Amendola, M. & van Steensel, B. Easy quantitative assessment of genome editing by sequence trace decomposition. Nucleic Acids Research 42, e168–e168 (2014).

71. Shalem, O. et al. Genome-Scale CRISPR-Cas9 Knockout Screening in Human Cells. Science 343, 84–87 (2014).

72. Livak, K.J. & Schmittgen, T.D. Analysis of Relative Gene Expression Data Using Real-Time Quantitative PCR and the 2−ΔΔCT Method. Methods 25, 402–408 (2001).

73. Wienholz, F. et al. FACT subunit Spt16 controls UVSSA recruitment to lesion-stalled RNA Pol II and stimulates TC-NER. Nucleic Acids Res 47, 4011–4025 (2019).

74. Tufegdžić Vidaković, A. et al. Regulation of the RNAPII Pool Is Integral to the DNA Damage Response. Cell 180, 1245–1261.e21 (2020).

75. Ramirez, F. et al. deepTools2: a next generation web server for deep-sequencing data analysis. Nucleic Acids Res 44, W160–5 (2016).

76. Gardner, J.M. & Jaspersen, S.L. Manipulating the yeast genome: deletion, mutation, and tagging by PCR. Methods Mol Biol 1205, 45–78 (2014).

77. Brachmann, C.B. et al. Designer deletion strains derived from Saccharomyces cerevisiae S288C: a useful set of strains and plasmids for PCR-mediated gene disruption and other applications. Yeast 14, 115–32 (1998).

78. Langmead, B. & Salzberg, S.L. Fast gapped-read alignment with Bowtie 2. Nat Methods 9, 357–9 (2012).

79. Mao, P., Smerdon, M.J., Roberts, S.A. & Wyrick, J.J. Asymmetric repair of UV damage in nucleosomes imposes a DNA strand polarity on somatic mutations in skin cancer. Genome Res 30, 12–21 (2020).

80. Park, D., Morris, A.R., Battenhouse, A. & Iyer, V.R. Simultaneous mapping of transcript ends at single-nucleotide resolution and identification of widespread promoter-associated non-coding RNA governed by TATA elements. Nucleic acids research 42, 3736–3749 (2014).

81. Eisen, M.B., Spellman, P.T., Brown, P.O. & Botstein, D. Cluster analysis and display of genome-wide expression patterns. Proceedings of the National Academy of Sciences 95, 14863 (1998).

82. Saldanha, A.J. Java Treeview--extensible visualization of microarray data. Bioinformatics 20, 3246–8 (2004).

83. Holstege, F.C. et al. Dissecting the regulatory circuitry of a eukaryotic genome. Cell 95, 717–28 (1998).

84. Mao, P. et al. Genome-wide maps of alkylation damage, repair, and mutagenesis in yeast reveal mechanisms of mutational heterogeneity. Genome Res 27, 1674–1684 (2017).

85. Weiner, A. et al. High-Resolution Chromatin Dynamics during a Yeast Stress Response. Molecular Cell 58, 371–386 (2015).

86. Hodges, A.J., Plummer, D.A. & Wyrick, J.J. NuA4 acetyltransferase is required for efficient nucleotide excision repair in yeast. DNA Repair (Amst) 73, 91–98 (2019).

87. Li, M., Ko, T. & Li, S. High-resolution Digital Mapping of N-Methylpurines in Human Cells Reveals Modulation of Their Induction and Repair by Nearest-neighbor Nucleotides. J Biol Chem 290, 23148–61 (2015).

88. Mukherjee, C. et al. RIF1 promotes replication fork protection and efficient restart to maintain genome stability. Nature Communications 10, 3287 (2019).

89. Callen, E. et al. ATM prevents the persistence and propagation of chromosome breaks in lymphocytes. Cell 130, 63–75 (2007).

90. Cornacchia, D. et al. Mouse Rif1 is a key regulator of the replication-timing programme in mammalian cells. Embo J 31, 3678–90 (2012).

91. van Cuijk, L. et al. SUMO and ubiquitin-dependent XPC exchange drives nucleotide excision repair. Nat Commun 6, 7499 (2015).

